# Mutators drive evolution of multi-resistance to antibiotics

**DOI:** 10.1101/643585

**Authors:** Danna R. Gifford, Ernesto Berríos-Caro, Christine Joerres, Marc Suñé, Jessica H. Forsyth, Anish Bhattacharyya, Tobias Galla, Christopher G. Knight

## Abstract

Antibiotic combination therapies are an approach used to counter the evolution of resistance; their purported benefit is they can stop the successive emergence of independent resistance mutations in the same genome. Here, we show that bacterial populations with ‘mutators’, organisms with defects in DNA repair, readily evolve resistance to combination antibiotic treatment when there is a delay in reaching inhibitory concentrations of antibiotic—under conditions where purely wild-type populations cannot. In populations subjected to combination treatment, we detected a remarkable amount of genomic diversity in resistance-determining mutations, multi-drug efflux pumps, and mutation-rate altering genes. However, using eco-evolutionary simulations, we demonstrate that only the initial mutator allele is required to explain multi-resistance evolution. Unexpectedly, mutators not only allowed multi-resistance to evolve under combination treatment where it was favoured, but also under single-drug treatments. Under both conditions, the mutator allele swept to fixation through hitch-hiking with single-drug resistance, enabling subsequent resistance mutations to emerge. Ultimately, our results suggest that mutators may hinder the utility of combination therapy when mutators are present. Additionally, by raising the rates of population mutation, selection for multi-resistance may have the unwanted side-effect of increasing the potential to evolve resistance to future antibiotic treatments.

**Significance statement:** The global rise in antimicrobial resistance means that we urgently need new approaches to halt its spread. Antibiotic combination therapy, treatment involving more than one antibiotic, is a strategy proposed to do just that. Evolving resistance to combinations is thought to be exceedingly rare, as it would require two independent mutations to occur in the same genetic background before microbial growth is inhibited. We find that wild-type populations cannot achieve this, even when antibiotic concentrations increase gradually. However, populations with ‘mutators’, organisms with elevated mutation rates through DNA repair defects, can readily evolve multi-drug resistance under both single-drug and combination treatments. Further, hitch-hiking of mutator alleles alongside resistance increases the evolutionary potential for acquiring further resistance mutations. As mutators are commonly found in natural populations, including infection, our results suggest that combination therapy may not be as resilient a strategy against resistance evolution as was once thought.

## Introduction

Rising rates of resistance and declines in antimicrobial discovery have lead to an emerging public health crisis. Consequently, there is an urgent need for strategies that suppress resistance evolution using existing antimicrobials. There has been sustained interest in the use of ‘combination therapy’ to prevent the development of antibiotic resistance in infectious diseases. ^1–3^ and cancer ^4^. Combination therapy uses multiple drugs as part of the same treatment, an approach that has proved successful in various settings ^5–9^. Much emphasis has been placed on characterising interactions between antibiotics i.e. ‘synergism’ and ‘antagonism’ ^3,10–12^, and how resistance mechanisms affect resistance to other antibiotics i.e. collateral sensitivity and cross resistance ^13–15^. There is significant interest in expanding the use of combination therapy to tackle the global burden of antimicrobial resistance ^16–18^. However, there is currently conflicting evidence that combining therapies can stop the emergence of resistance, with some combinations performing no better than monotherapy in some contexts ^19,20^. Determining what governs the resilience of combinations against resistance evolution therefore remains an open question.

The ability for combination therapy to prevent resistance is predicated on the requirement for multiple independent resistance mutations to achieve multi-drug resistance. Due to low rates of spontaneous mutation for various antibiotics ^21^, obtaining multiple independent mutations before growth inhibition is achieved assumed to be exceedingly rare ^1,22^, discounting multi-drug resistance and cross-resistance mechanisms ^23^. Simultaneous inhibition caused by two drugs prevents independent mutations from occurring sequentially, and resistance typically does not spread in the absence of selection. However, this assumes inhibition is achieved before resistance can emerge, which due to drug pharmacokinetics may be unrealistic. Exposure to sub-inhibitory concentrations may enable the spread of single-resistance mutations ^24–27^.

In contrast to the characterisation of drug interactions, the effect of variation in microbial populations on the resilience of combination therapies has received far less attention. Specifically, natural bacterial populations frequently contain ‘mutators’, variants with defects in DNA replication and repair. Mutation rates of mutators are typically 10-to 1000-fold higher than wild-type ^28,29^. Mutator frequencies can be as high as 30% e.g. in infections and host-associated microbiomes ^30–33^. High rates of resistance are often found in mutator lineages ^28,34–37^, and current support for the efficacy of combinations against mutators is mixed ^38–40^. Understanding the evolutionary dynamics of mutators and multi-resistance evolution is therefore important for predicting whether combination antibiotic therapies may be effective at preventing resistance.

In this study, we assessed the evolutionary dynamics of multi-resistance in populations with mutators at initial frequencies found in natural populations (0-30%). Using a combination of experimental evolution and stochastic modelling, we challenged populations of *Escherichia coli* with evolving antibiotic resistance to rifampicin and nalidixic acid, either used singly or in combination. We imposed selection for resistance using a ‘ramping selection’ design over physiologically-relevant concentrations of antibiotics with a delay in reaching inhibition ^41,42^. This enables us to mimic environmental variation akin to what occurs in clinical (e.g. within-patient pharmacokinetics ^26^) and environmental settings (e.g. wastewater contamination ^43^, and agriculture ^44^). This approach provides the opportunity to test directed hypotheses regarding the role of mutators due to a solid understanding of the underlying evolutionary processes involved in resistance to these antibiotics.

Our results demonstrate that sub-populations of mutators facilitate the evolution of multi-resistance. Multi-resistance arose by independent single-resistance mutations in populations where mutators are present, but not in purely wild-type populations. Genome sequencing revealed that multi-resistance arose due to mutations in the targets of rifampicin ^45^ and nalidixic acid ^46^. We also observed variation across the genome, including, for some isolates, mutations in multi-drug efflux systems and DNA replication and repair systems. However, stochastic simulations revealed that differences in the rate of mutation at the two canonical targets, allowing resistance to evolve sequentially, was sufficient to explain multi-resistance evolution in these populations. Multi-resistance was able to evolve without requiring multi-drug resistance mechanisms, cross-resistance, spontaneous double mutations, a further increase in mutation rate or any other genomic variation. Strikingly, multi-resistance arose both under combination and single-drug treatments, due to hitch-hiking of the mutator allele alongside single-drug resistance leading to an increase in mutation rate. Given the prevalence of mutators in natural populations, multi-resistance evolution may represent a considerable challenge to the wider deployment of combination therapy against bacterial infections. Moreover, because multi-resistance emerged in both single-drug and combination treatments, genetic variation underlying resistance to combination treatments may already be present, even before the use of such treatments becomes common practice.

## Results

### Multi-resistance evolves in both single-drug and combination treatments when mutators are present

To determine the conditions under which multi-resistance evolves, we then performed experimental evolution using four mutator frequency treatments (none, low, intermediate, high) and four selection regimes (antibiotic-free, single-drug with either rifampicin or nalidixic acid, combination with both antibiotics). We employed ramping selection, where antibiotic concentrations were doubled daily over six days (from 0.625 mg/l to 20 mg/l of each drug, where 10 mg/l of either is sufficient to inhibit wild-type growth). Figure 1 shows the number of populations with resistance in each treatment. Multi-resistance evolution was not observed in the absence of mutators or in the absence of selection for resistance (although single resistance was observed). However, when mutators were present, multi-resistance evolved in both single-drug treatments and the combination treatment. While the combination treatment was most effective overall at suppressing resistance, a considerable proportion of populations with mutators (23.3%–61.7%) nevertheless evolved multi-resistance.

**Figure 1:**
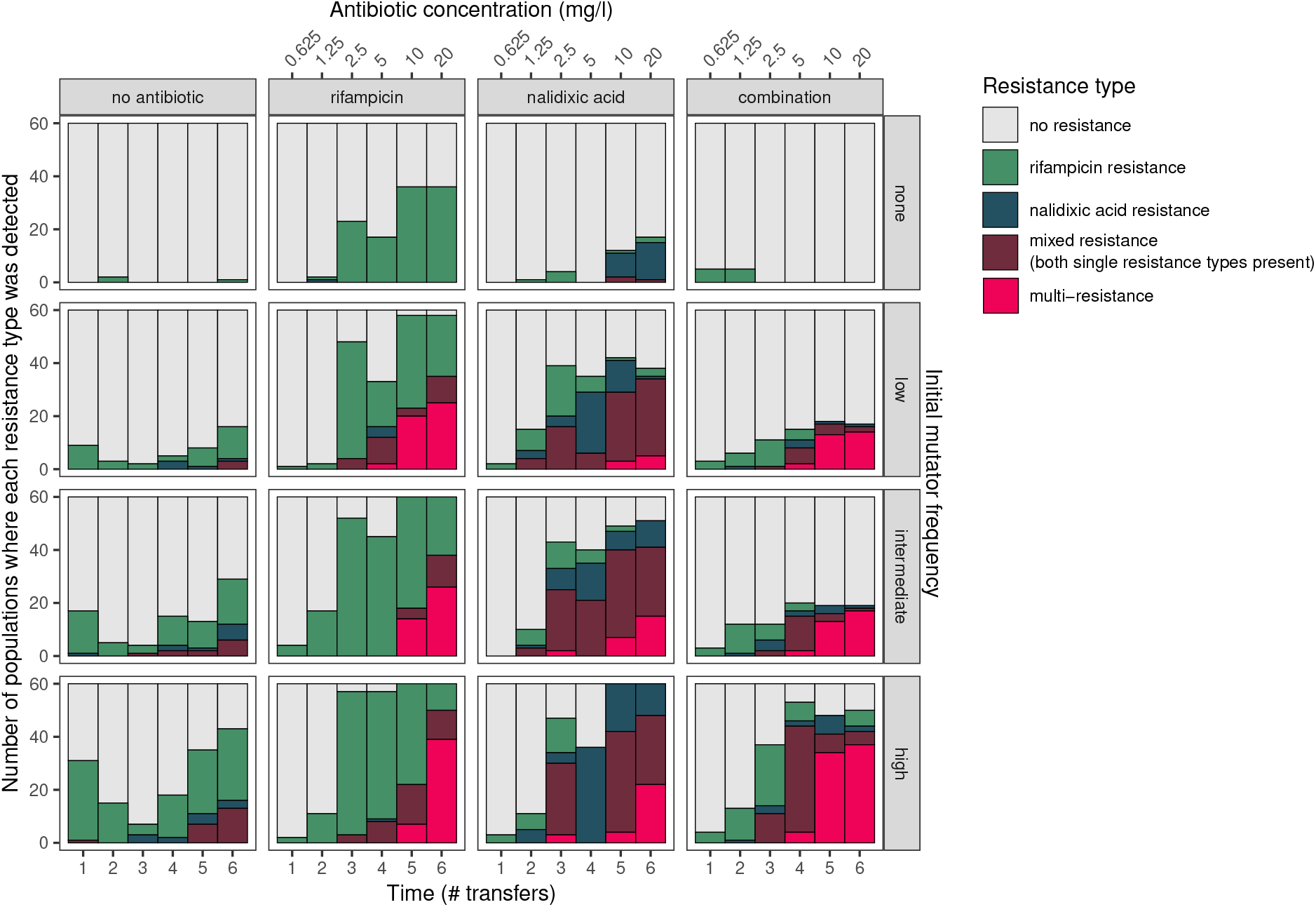
Multi-resistance evolves in populations where mutators are present in both single-drug and combination drug treatments. ‘Resistance type’ is characterised by growth on single-drug (‘rifampicin resistance’, ‘nalidixic acid resistance’) or combination (‘multi-resistance’) selective media; ‘mixed resistance’ refers to growth on both single-drug media, but not combination selective media.

The effect of mutator initial frequency and antibiotic treatment on resistance state of populations at the end of the experiment was determined using a Bayesian categorical mixed-effects model (see Supplementary Table 1, Supplementary Figure 1, and associated text). The presence of any frequency of mutators increased the proportion of populations exhibiting single-drug resistance (including mixed resistance, where the population contained organisms capable of growing in either single-drug environment, but not the two-drug environment) or multi-resistance. There was a positive association between initial mutator frequency and resistance, although with overlapping 95% C.I.s.

### Phenotypic and genomic characterisation of evolved multi-resistant isolates arising during ramping selection experiment

We assayed growth phenotypes and performed whole genome sequencing on evolved multi-resistant isolates. There was a strong correlation between growth in minimum and maximum concentrations of the combination treatment (i.e. 0 mg/l and 20 mg/l) (Figure 2A, Bayesian multivariate regression: *r* = 0.73, 95% C.I.: (0.57, 0.84), see Supplementary Figure 2, Supplementary Table 2). There was little effect of mutator frequency on growth, although there was a tendency toward lower growth in isolates from the ‘low’ frequency treatment.

**Figure 2:**
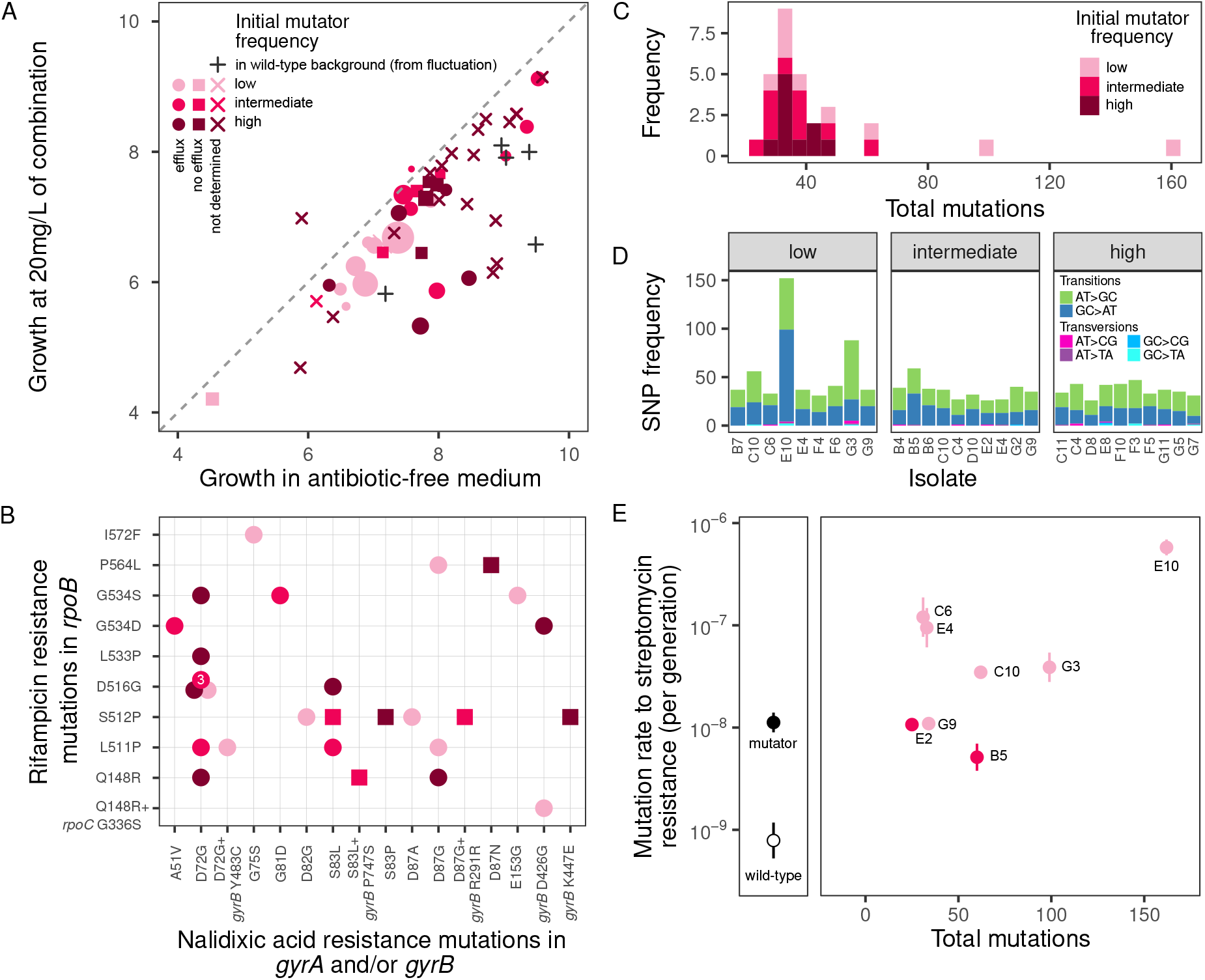
Phenotypic and genomic changes underpinning multi-resistance evolution in mutators. A. Growth of multi-resistant mutants under initial and final conditions of the selection experiment (i.e. 0 mg/l and 20 mg/l of the combination; growth measured as AUC of OD 600 nm curves over 22 h; *n* = 50). For sequenced isolates (*n* = 29), point shape indicates isolates with (‘circles’) or without (‘squares’) efflux pump mutations, and point size reflects total number of acquired mutations. B. Mutations identified in canonical drug resistance targets for rifampicin and nalidixic acid resistance (shapes and colours are as in panel A). C. Number of mutations identified in each sequenced multi-resistant isolate. D. Spectrum of spontaneous SNPs is dominated by transitions. E. Number of mutations detected by whole genome sequencing is positively associated with mutation rate in multi-resistant isolates (as measured by fluctuation test to streptomycin resistance, *n* = 8).

We performed whole genome sequencing and variant calling on a subset of multi-resistant isolates (Supplementary Table 3 and Supplementary File 1). All isolates acquired rifampicin resistance mutations in the canonical mutational target *rpoB*, with one strain acquiring an additional mutation in *rpoC*; for nalidixic acid resistance, 23/29 possessed a single mutation in *gyrA*, 2/29 a single mutation in *gyrB*, and 3/29 acquired one mutation in each of *gyrA* and *gyrB*. Diverse SNPs were observed in both of these targets. There was a significant association between specific SNPs occurring in *rpoB* and *gyrA* (Fisher’s exact test, *p* = 0.04), which arose because the pair *rpoB* D516G and *gyrA* D72G occurred 5 times in total (Figure 2B). Most strains (23/29) also acquired mutations in multi-drug efflux pump genes, the most frequent being *acrR* (9/29), a repressor involved in the AcrAB-TolC system ^47^. However, efflux pump mutations included both synonymous substitutions and putative loss-of-function mutations in structural components, which are unlikely to improve resistance.

In addition to resistance-associated mutations, we detected mutations across the genome (median = 34, range = 25–162, Figure 2C). The mutational spectrum of SNPs was dominated by transitions (Figure 2D), which is characteristic of the specific Δ*mutS* mutator allele introduced here ^48,49^. While some of this additional variation likely affected fitness, there was little association between the number of mutations identified through whole genome sequencing and growth (Figure 2A). Many mutations observed were likely to be selectively neutral, as 283/1252 (22.6%) of all mutations detected were synonymous SNPs. Notably, we identified exceptionally high numbers of mutations in 4/29 isolates (three isolates with 62, 99, and 162 mutations each from the ‘low’ treatment, and one isolate with 60 mutations from the ‘intermediate’ treatment). We therefore assessed whether mutation rate itself had evolved during the experiment by measuring the mutation rates of eight multi-resistant isolates (the four mentioned above and four matched isolates from the same treatments), in comparison with the original Δ*mutS* strain and the wild-type. There was a positive association between estimated mutation rate and number of mutations each isolate had acquired during selection (Figure 2E). While three evolved isolates had estimated mutation rates approximately equal to (or slightly lower than) Δ*mutS*, five had increased mutation rates, ranging from 3-fold to 51-fold greater than Δ*mutS*—among most of these, we detected mutations in DNA replication and repair genes (Supplementary Table 4).

### Stochastic simulations reveal key mechanisms of multi-resistance evolution

To gain insight into the drivers of multi-resistance evolution, including the requirement for different classes of mutations observed, we used a stochastic population-dynamic simulation model. We produced a minimal model capable of predicting multi-resistance evolution, i.e. we included only a single allele for each resistance type, and excluded multi-drug resistance mechanisms, spontaneous double mutants, recombination, and mutations at non-resistance-conferring loci. Disallowing these types of variation permitted us to test hypotheses regarding the roles of selection and mutation supply on the emergence of multi-resistance. Populations initially contained only sensitive individuals, which we denote *S* for the wild-type (and *S*′ for the mutator, if present, differing only in mutation rate from the wild-type). During growth, single mutations can occur, giving rise to rifampicin-resistant type *R* (and *R*′) and nalidixic acid resistant type *N* (and *N*′). Subsequently, *R* and *N* (and *R*′ and *N*′) can each give rise to multi-resistant type *D* (and *D*′). We used a stochastic approach because it more accurately captures the dynamics of mutations that arise in a single cell, i.e. they occur as random events ^50–52^. For each set of conditions, we simulated 1000 populations with a maximum population size of 5.71 × 10^8^ bacteria, equivalent to the maximum density observed in the selection experiment. Simulation parameters relating to growth were estimated empirically from growth curve data using strains that were derived from fluctuation tests (i.e. independent from the selection experiment, Supplementary Figure 6; see Supplementary Information for full details). We estimated mutation rates of our wild-type and mutator strains in previous work ^21,53^.

This simulation model captured the major features of the wet-lab experiments (Figure 3, cf. Figure 1), i.e. that multi-resistance was constrained to populations treated with antibiotics, including single antibiotic treatments, and that the presence of mutators facilitated multi-resistance evolution. A Bayesian categorical model fitted to these simulations produced parameter estimates that closely matched those from the experimental data, demonstrating that the simulations quantitatively recapitulate the experiments (Supplementary Figure 10). A small fraction of purely wild-type simulated populations evolved multi-resistance (between 0/1000 and 15/1000, depending on treatment); this is consistent with expectations from our experimental results (< 1/60 per treatment). As the simulation excludes the possibility of multi-resistance emerging through a single reproductive event, this suggests that multi-resistance can emerge without invoking multi-drug resistance mechanisms (e.g. efflux pumps), simultaneous acquisition of two resistance mutations, or recombination.

**Figure 3:**
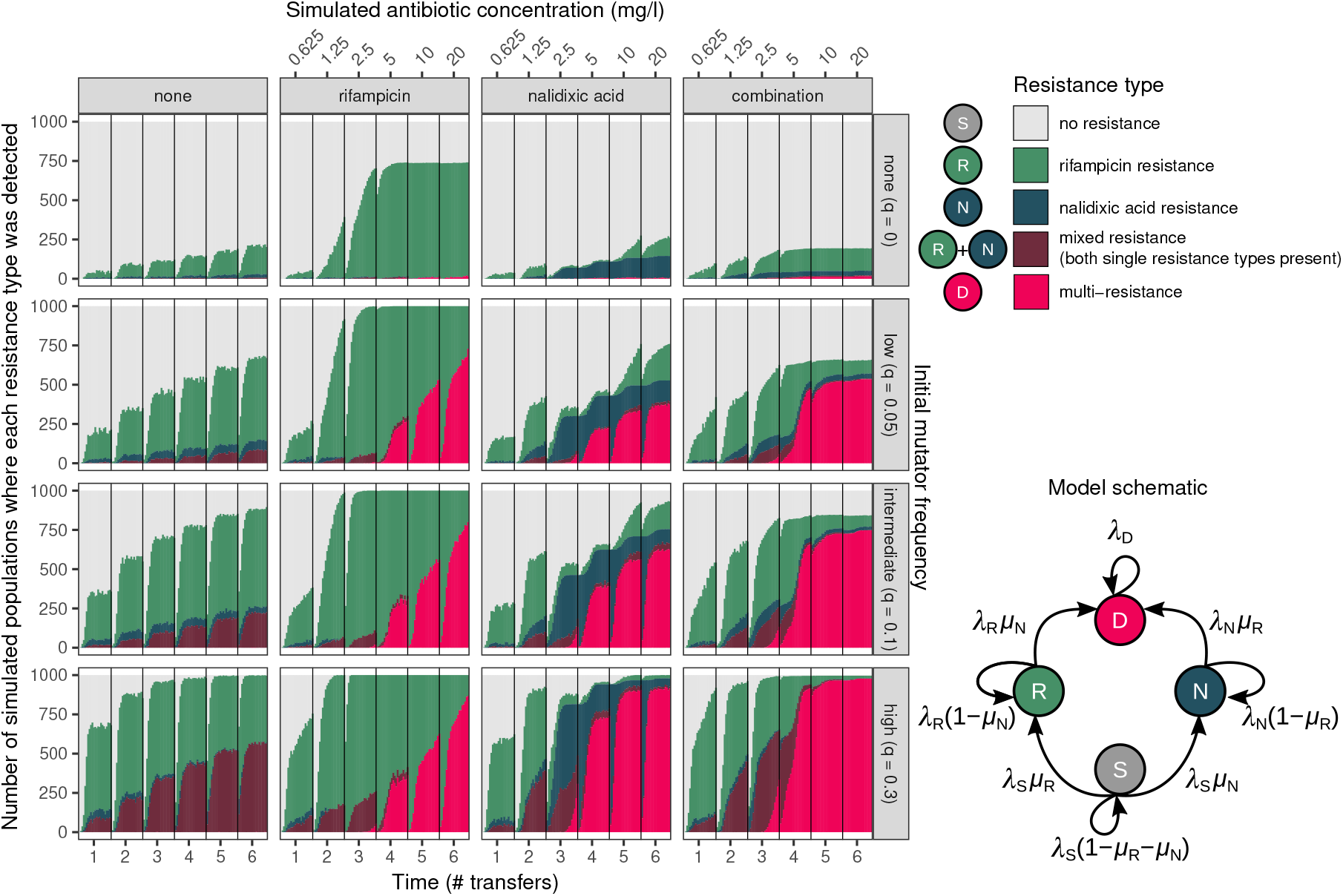
Simulated resistance evolution allowing only sequential acquisition of resistance recapitulated the emergence of resistance in experiments, suggesting elevated single-drug resistance mutation rates are sufficient to explain multi-resistance evolution. Resistance type was determined in an equivalent manner to experimental data in Figure 1 (i.e. if at least on individual of each type was detected in a random sample of 1/200 of each population, see Methods). The schematic depits the the simulation approach, where *λ*_*i*_ refers to the reproduction rate of Type *i*, and *µ*_*j*_ refers to the mutation rate to resist antibiotic *j* (see Supplementary Information for full details). Parameters used in simulations are shown in Supplementary Figure 6.

Simulations enabled us to observe the dynamics of multi-resistance evolution within each population. Figure 4A shows the most commonly observed evolutionary trajectory to multi-resistance, *S*′ → *R*′ → *D*′; examples of other trajectories can be found in Supplementary Figure 14. In general, single resistance arose early in the mutator genetic background, but remained at low frequency until it conferred a fitness benefit over the sensitive type. Once the single-resistant lineage began to increase in frequency, a subsequent mutation producing the multi-resistant type emerged. In some cases, resistant types can be lost due to population bottlenecks (as is the case for *N*′ in Figure 4A and *R*′ and *D*′ in Supplementary Figure 14); these may re-emerge and go on to found a *D*′ type lineage, or ultimately go extinct. *R*′ and *N*′ types can sometimes emerge and co-exit without going extinct, ultimately leading to two independent lineages resulting in *D*′ type individuals. These within-population dynamics are summarised across all replicate simulations Figure 4B (see also Supplementary Figure 15). This demonstrates that, while selective sweeps for single-resistance were observed in all antibiotic-containing treatments, multi-resistance only swept to fixation in the combination treatment.

**Figure 4:**
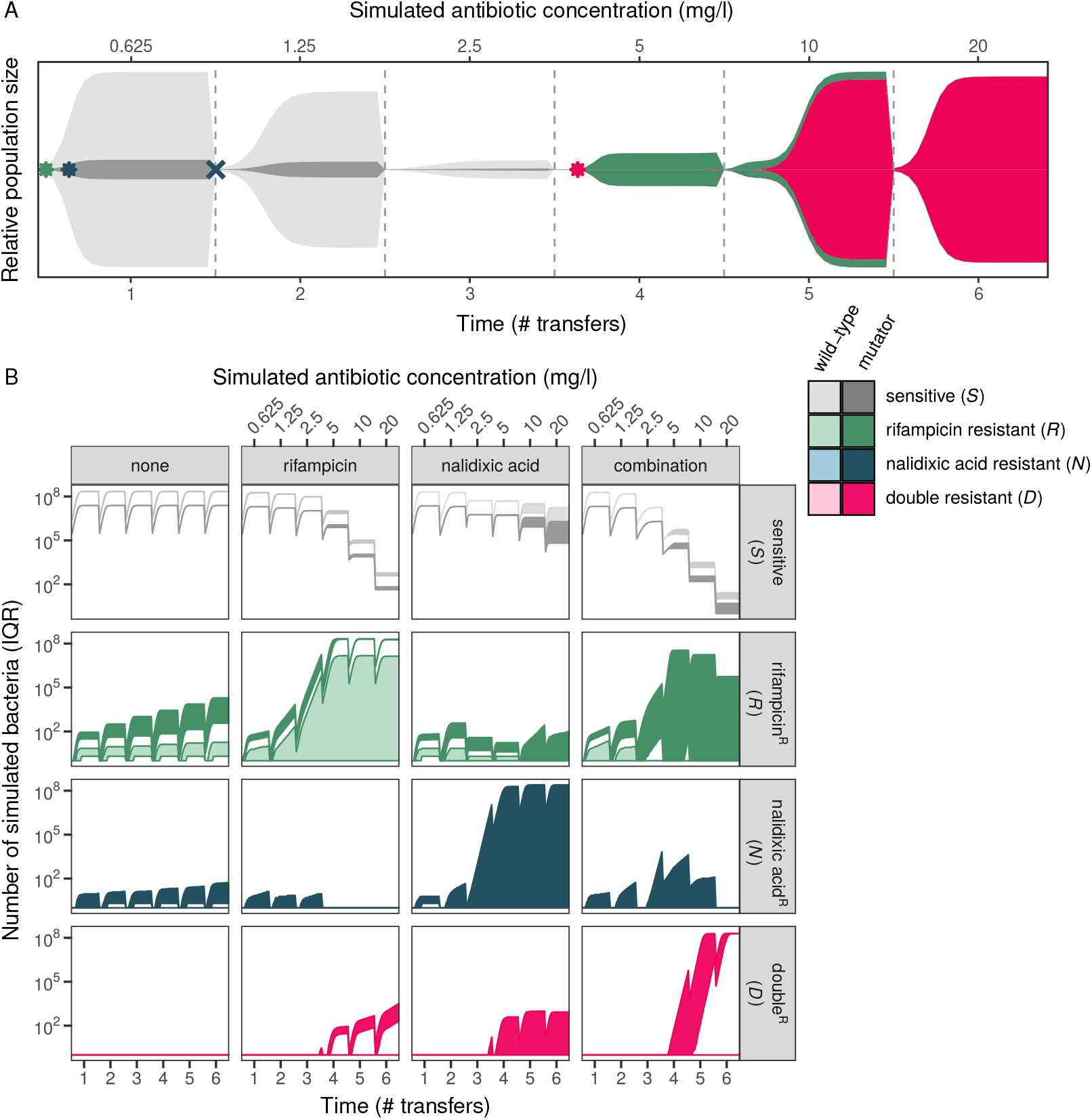
Population dynamics of simulated resistance evolution. A. Muller diagram showing typical progression from sensitive to multi-resistant in the combination treatment within a single population. Multi-resistance most commonly emerged via *S*′ → *R*′ → *D*′ in the mutator genetic background (examples of other paths are shown in Supplementary Figure 14). Height of each area corresponds to the population size of each type. Example shown is a single replicate from the ‘intermediate’ initial mutator frequency (*u* = 0.1) treatment. B. Number of bacteria of each resistant type for the four simulated treatments (interquartile range, IQR, over *n* = 1000 replicate simulations). Results shown from the ‘intermediate’ initial mutator frequency (*u* = 0.1; other frequencies are shown in Supplementary Figure 15).

To evaluate the generality of these findings with regard to mutators, we ran further simulations with different parameter values. We first explored different dose escalation schemes, varying the amount of time until MIC is reached. We find that mutators facilitate multi-resistance evolution when the time needed to reach MIC decreases four-fold or increases five-fold (see Supplementary Figure 11 and associated text). We also assessed whether these findings are specific to the particular empirical growth curves resulting from exposure to rifampicin and/or nalidixic acid. We modified the simulations to consider simple logistic growth with an imposed ‘cost of resistance’ for single- and multi-resistance, as such costs may affect the spread of multi-resistance ^54^, but *cf*. ^55^). We observed the same general patterns of multi-resistance with imposed fitness costs as obtained with our empirical parameter estimates (see Supplementary Figure 12 and associated text). Finally, we explored whether large population sizes could allow multi-resistance evolution in the absence of mutators. We found that, in the combination treatment, population sizes > 5.71 × 10^10^ evolve multi-resistance in the absence of mutators at an appreciable rate (Supplementary Figure 13), although this represents a population size much larger than most typical infections ^56–59^.

## Discussion

A major motivation behind using antibiotic combination therapy is its presumed resilience against resistance evolution. However, our results question this resilience under two common scenarios that occur in clinic and nature: delayed inhibition and the presence of mutators. Multi-resistance evolved under both single-drug and combination treatments when mutators were introduced at frequencies often found in infection (Figure 1), despite conferring no clear advantage in single-drug environments (Supplementary Figure 6). Increased mutation rate brought on by defective mismatch repair was sufficient to explain the emergence of multi-resistance via sequential acquisition of independent resistance mutations. Although other evolutionary mechanisms could be relevant in other contexts, they need not be invoked here, e.g. differences in birth and death rates affecting the emergence of resistance ^60^, acquiring fitter resistance alleles through clonal interference ^61^, differential supply of compensatory mutations ^62^, genomic variation influencing MIC ^63^, or co-evolution between genome and resistance mechanisms ^64^. Our findings raise concerns about the effectiveness of combination treatment in combating the evolution of drug resistance. Due consideration should be given to the evolutionary consequences of combination therapy as mutators are frequently present in infections—particularly in chronic infections, for which combination therapies are especially important.

To reveal the key evolutionary mechanisms that allow mutators to evolve multi-resistance, we first characterised the phenotypic and genomic changes that occurred in isolates that evolved resistance to both antibiotics (Figure 2). All sequenced isolates had acquired mutations in the drug targets of rifampicin (*rpoB*) and nalidixic acid (*gyrA* and/or *gyrB*). Most isolates also acquired mutations in multi-drug efflux pumps, consistent with findings in other organisms ^65^. However, efflux pump mutations were not present in all isolates, and were never found without canonical single-drug resistance mutations, indicating that multi-drug resistance mechanisms were neither sufficient nor required to achieve multi-resistance here. Isolates also possessed remarkable genomic diversity without any direct association with resistance. Much of this variation is likely selectively neutral as there was no association between numbers of mutations acquired and fitness (antibiotic-free: *r* = −0.01, combination: *r* = −0.06, Figure 2) and many observed mutations were synonymous SNPs. However, there is evidence that mutation rates themselves evolved. For some isolates, we found a 3-to 51-fold increase in mutation rate relative to the ancestral f..*mutS* strain. We found mutations in genes associated with DNA replication and repair, which likely hitch-hiked alongside multi-resistance as it swept toward fixation. Selection for multi-resistance via mutator alleles therefore appears to come at little cost to the organism, while also sometimes modifying genes associated with multi-drug resistance (efflux pumps) and increasing the evolutionary potential for developing resistance to other antibiotics (via selective sweeps of mutators and/or mutation rate evolution). These unintended consequences of combination therapy are alarming, and suggest that thoughtful consideration of the population genetics of multi-resistance need consideration before combination therapies are deployed more broadly.

To ascertain which forms of genetic variation were essential for multi-resistance evolution, we carried out *in silico* stochastic simulations to determine whether multi-resistance could arise via acquiring two independent resistance mutations alone. The simulations closely matched the experimental results, suggesting that, in spite of other variation observed, the original mutator allele was sufficient to enable multi-resistance evolution. This provides further evidence that the multi-drug efflux pump alterations and mutation rate increases seen in the selection experiment were not required for multi-resistance. In addition, the simulations also allowed us to delve into the evolutionary dynamics of multi-resistance occurring in individual populations. In all cases, mutator alleles hitch-hiked alongside single-drug resistance mutations, which then permitted a subsequent resistance mutation to occur in the same genetic background consistent with previous work ^66–69^. However, different antibiotic treatments had large differences in the frequency of multi-resistant individuals within the population: in the combination treatment, multi-resistance swept toward fixation, but achieved only low frequency in single-drug treatments where it confers little-to-no fitness benefit (Figure 4).

Mutators present other challenges for infection management beyond increasing the propensity for resistance, e.g. mitigating fitness costs of antibiotic resistance through compensatory adaptation ^62^, gaining resistance to non-antibiotic forms of bacterial control, such as vaccination ^70^ and phage therapy ^71^, and adapting to host conditions in opportunistic pathogens ^72^. Mutators are also unlikely to suffer from diminished fitness due to the accumulation of deleterious mutations (i.e. ‘lethal mutagenesis’ or ‘mutational meltdown’ ^73–75^), as we found no evidence of reduced fitness in highly-mutated isolates (Figure 2A), consistent with previous short-term ^76^ and long-term experiments ^77^ excepting experiments where populations are kept artificially small, ^78^. Together, this suggests that, once multi-resistant mutator lineages become established, they will be difficult to eradicate—whether through natural selection or through alternatives to antibiotics. This raises the possibility that screening for mutators, in addition to antibiotic susceptibility, could be valuable in clinical practice.

Although our emphasis here was on mutators, our results are likely applicable to other factors that affect mutation rates, e.g. environmental factors such as nutrients and temperature ^53,79–82^, stress-induced mutagenesis ^83–86^, radical-induced DNA damage ^87,88^, and biotic interactions ^21,89–91^. Theoretical models have also predicted that increases in mutation rate *variability*, not just average, will lead to a higher probability of evolving multi-resistance ^92^. A better understanding of these varied influences on mutation rates will be critical in the application of methods for preventing resistance through reducing the supply of resistance mutations ^93–95^.

A vast number of combinations can be generated from existing antibiotics ^96^, which makes combination therapy an enticing approach for countering the rise of drug-resistant infections. While combinations treatments are indeed useful for reducing resistance evolution ^38,39^, our results suggest that the potential for multi-resistance evolution needs thoughtful consideration in the design and application of such treatments. As direct assessment of all possible combinations is likely an insurmountable task ^96^, the experimental and modelling approaches developed here can serve as a framework for predicting whether particular combinations can suppress resistance evolution.

## Acknowledgements

This project was supported by the BBSRC (DRG and CGK: BB/M020975/1, CJ: BB/M011208/1), a UKRI Innovation/Rutherford Fund Fellowship (DRG: MR/R024936/1), the Academy of Medical Sciences (DG: SBF007\100096), a University of Manchester Presidential Scholarship (EBC), a Postdoctoral Seed Award from Earth and Environmental Sciences, The University of Manchester (DRG), and a Wellcome Trust Institutional Strategic Support Fund award (DRG, MS, TG and CGK: part of 204796/Z/16/Z). TG acknowledges support from The Maria de Maeztu program for Units of Excellence in R&D (MDM-2017-0711). MS acknowledges the support of the Swedish Research Council (Grant No. 638-2013-9243). The funders had no role in study design, data collection and analysis, preparation of, or decision to publish the manuscript. The authors thank A Wilkinson for access to warm-room facilities, and N Cochrane for guidance on bacterial population sizes relevant to infection. The authors acknowledge assistance from Research IT and the Computational Shared Facility at The University of Manchester. DRG wishes to thank TD Dubé for a discussion on combination therapy that inspired this work.

## Author contributions

Conceived of experiment: DRG, CGK. Conducted experiments: CJ, JF, DRG. Analysis of experimental data: EBC, DRG. Stochastic simulations: EBC, MS, DRG, TG. Genome sequencing and analysis: AB, DRG. Initial draft of manuscript: DRG, EBC, CGK. Edited and approved final manuscript: all authors.

## Data availability

Data, R scripts, and source code are available on GitHub (https://github.com/dannagifford/multi-resistance/).

## Methods

### Strains and media

Selection experiments involved ‘wild-type’ *E. coli* str. K-12 substr. BW25113 [F-, Δ (*araD*-*araB*)567, Δ*lacZ* 4787(::*rrnB*-3), *λ*-, *rph*-1, Δ (*rhaD*-*rhaB*)568, *hsdR*514] ^97^, and a ‘mutator’ strain Δ*mutS* (as above, but with Δ*mutS*738::kan, indicating Δ*mutS* replacement with kanamycin resistance). The kanamycin resistance cassette has not previously been observed to affect resistance to the antibiotics we have considered here ^21,53^. Both strains were obtained from Dharmacon, Horizon Discovery Group, UK. Relative to the published reference genome ^98^, whole genome resequencing revealed no pre-existing mutations in the wild-type BW25113 background, and a single point mutation in the f..*mutS* strain (1,985,889 G>A, resulting in an amino acid substitution in *pgsA* A137V), which does not have a known association with resistance.

Routine culturing was performed in lysogeny broth [LB, 10 g/l tryptone (Fisher Scientific, UK), 5 g/l Bacto yeast extract (BD Biosciences, UK), 10 g/l NaCl (Fisher Scientific, UK)]. Selection experiments in the presence of antibiotic(s) were performed in Müller-Hinton broth (MH broth, 23 g/l, Sigma-Aldrich, UK). MH is a preferred medium for use with antibiotics ^99^. Solid media were made by adding 12 g/l agar (BD Biosciences, UK) to either broth prior to autoclaving. Stock antibiotic solutions were prepared at 10 mg/ml. Rifampicin (Fisher Scientific, UK) was dissolved in methanol (Fisher Scientific, UK), and nalidixic acid (Fisher Scientific, UK) was dissolved in double distilled water, with 1N NaOH (Fisher Scientific, UK) added drop-wise until the antibiotic was solubilised. Strains were stored in LB with 40% glycerol at −80 °C.

### Selection experiment under single-drug and combination treatments

We used experimental evolution to determine the effect of mutators on multi-resistance evolution under single and combination antibiotic treatments. Populations were founded from a mixture of mutator and wild-type individuals. Independent overnight cultures of wild-type and mutator were first grown separately in 5 ml MH broth. Volumetric mixtures of the mutator and wild-type overnight cultures were made at ratios of 0/100% (‘none’), 10/90% (‘low’), 25%/75% (‘intermediate’), and 50%/50% (‘high’) reflecting mutator frequencies observed in host-associated populations ^30–33^. We measured the actual frequency of mutators (denoted *u*) by plating serial dilutions of the populations on LB with 100 mg/l kanamycin agar (mutator count) and on LB agar (total population count). The initial mixtures were assayed for resistance to rifampicin or nalidixic acid by plating on MH agar supplemented with rifampicin (50 mg/l) or nalidixic acid (30 mg/l). Five cultures were discarded for having detectable rifampicin resistance at the beginning of the experiment; no cultures had detectable nalidixic acid resistance.

We used a serial transfer protocol that exposed populations to increasing concentrations of antibiotics over a period of six days. The experiment was performed in 96-well microtitre plates (Nunc, Fisher Scientific, UK) in 200 µl volumes grown at 37 °C with 200 rpm shaking in an Innova 42R Incubator (Eppendorf, United Kingdom) for 22 h growth periods (‘days’). Position of each antibiotic treatment within the plate was assigned using stratified randomization. Populations were initiated from the mixed cultures by diluting 1 µl of each into 200 µl of fresh culture using a 96-pin replicator (Boekel Scientific, Feasterville, PA, USA). At the end of each day, 1 µl of each population was pin replicated into 200 µl of fresh growth medium. This experimental protocol allows for a maximum of ∼ 46 generations of evolution (i.e. six days × 7.64 doublings/day). However, in practice the number of generations achieved by non-resistant strains in the presence of antibiotic(s) is likely to be less, given reduced carrying capacity at higher antibiotic concentrations (see Supplementary Figure 6).

Four antibiotic treatment regimes were used: no antibiotic, rifampicin only, nalidixic acid only, or rifampicin and nalidixic acid combined. Antibiotic concentrations were doubled each day over the course of six days (0.625, 1.25, 2.5, 5, 10, 20 mg/l of each individual antibiotic), where inhibition of the ancestral strain was achieved on day 5 a protocol often used in resistance evolution experiments, ^3,100–102^. The concentrations reflect physiological concentrations achieved during treatment with rifampicin maximum serum concentration 5-18 mg/l, ^103^ and nalidixic acid maximum plasma concentration 1.8-30 mg/l, ^41^. We note that nalidixic acid itself is mutagenic approximately two-fold increase at 10 mg/l ^104^, but the effect is minor relative to Δ*mutS* deletion.

### Detection and analysis of resistance

Following each transfer, we used a high-throughput resistance assay involving pin replicating 1 µl of overnight culture (equivalent to a random sample of 1/200th of the population) on MH agar without antibiotic, or with rifampicin (50 mg/l), nalidixic acid (30 mg/l), or rifampicin and nalidixic acid combined (50 mg/l and 30 mg/l, respectively) in 120 mm square Petri dishes. Growth at these concentrations is indicitave of mutations in the canonical resistance genes for these antibiotics in *E. coli, rpoB* and *gyrA*, respectively. Populations were determined to be one of five ‘resistance states’: ‘sensitive’ (growth only on non-selective plates), ‘rifampicin resistant’ or ‘nalidixic acid resistant’ (growth only one of the two single-drug plates), ‘mixed resistant’ (growth on both single-drug selective plates but not combination selective plates), and ‘multi-resistant’ (growth on combination selective plates, as well as single-drug selective plates). Note these outcomes refer to *establishment* of resistance, rather than *fixation*, i.e. the proportion of resistant individuals is *>* 0 and ≤ 1. We analysed the probability of observing each resistance type using a Bayesian categorical model, implemented in the `brms` package ^105,106^ in R 3.5.3^107^, described in full in the Supplementary Information.

### Growth parameters of single- and multi-resistant clones

To determine the effects of single and multi-resistance on growth parameters, we selected five nalidixic acid resistant and five rifampicin resistant clones arising from the wild-type BW25113 genetic background via fluctuation tests ^108^, using an established protocol ^109^. Briefly, 1 ml LB cultures of *E. coli* K-12 BW25113 were grown overnight in 96-well deep-well plates. The entire volume of each culture was plated on MH agar supplemented with rifampicin (50 mg/l) or nalidixic acid (30 mg/l) in the wells of a 6-well plate (each well approximately 35 mm in diameter). These 6-well plates were incubated for 48 h. To select multi-resistant clones, we performed a second fluctuation test using resistant strains from the first, plating on the antibiotic to which they were not already resistant. Colonies were isolated from selective plates, grown overnight in LB medium, and then stored at −80 °C.

Growth curves were generated by measuring optical density (OD) at 600 nm every 30 min for 45 h using a BMG FLUOstar OMEGA with Microplate Stacker (BMG Labtech, Ortenberg, Germany, Supplementary Figure 3A). Each clone was grown in duplicate at 37 °C under each of the antibiotic concentrations experienced during the selection experiments (i.e. 0.625, 1.25, 2.5, 5, 10, 20 mg/l each of rifampicin and/or nalidixic acid). Cultures were initiated by first growing clones overnight in 200 µl MH broth, then diluted 1/200 into a total volume of 200 µl MH broth containing one or both antibiotic(s). The growth curves were used to estimate parameters for the stochastic simulation model using a custom MATLAB script (see ‘Data availability’ statement). The fitting procedure is described in full the Supplementary Information.

In addition to fitting growth curves, we also summarised growth curves into a single metric, area under the curve (AUC, Supplementary Figure 3B), as the empirical growth curves did not follow a standard logistic shape (which we also account for in the simulations below). We calculated AUC using the SummarizeGrowth function from the R package growthcurver ^110^. AUC incorporates all of lag phase, growth rate, and density, and highly repeatable. To determine whether multi-resistance conferred a benefit under single-drug treatments, we fit a Bayesian multivariate linear regression model of AUC of different strains in the presence of each treatment over all concentrations (described in full in the Supplementary Information).

Using the same protocol as above, growth curves for the multi-resistant clones that evolved during the selection experiment (all in the mutator genetic background) were measured in antibiotic-free medium, and in 20 mg/l of the combination treatment. The association between AUC, initial mutator frequency and treatment was analysed using a Bayesian multivariate regression model (described in full in the Supplementary Information).

### Mutation rate estimates and the probability of spontaneous double mutants

Mutation rates to rifampicin resistance (*µ*_*R*_ = 6.7 × 10^−9^ per cell division) and nalidixic acid resistance (*µ*_*N*_ = 7.4 × 10^−10^ per cell division), were obtained for the these strains in a previous publication ^53^, as was the mutator effect of Δ*mutS* (80-fold increase relative to wild type) ^21^. Assuming mutations occur independently, we estimate the probability of both rifampicin and nalidixic acid resistance mutations occurring simultaneously during the same replication event (i.e. a ‘spontaneous double mutant’) for the wild-type as the product of their mutation rates to resistance ^1,22^, i.e. *µ*_*R*_*µ*_*N*_ = 5.0 × 10^−18^, and for mutators as *µ*_*R*_*µ*_*N*_ × 80^2^ = 3.2 × 10^−14^. We can use these probabilities to obtain a rough estimate for the probability of observing a spontaneous double mutant by simulating a fluctuation test using `rflan()` from the R package ‘flan’ ^111^. We performed 10^6^ simulations using the same parameters as the selection experiment (i.e. 60 independent populations, days 1–4 permitting wild-type growth, maximum population size of 5.71 × 10^8^). For populations comprising only wild-type individuals, no spontaneous double mutants were observed in 10^6^ simulations. For populations comprising solely mutators, a spontaneous double mutant was observed in 4210 out of 10^6^ simulations. Hence, the probability of having observed a spontaneous double mutant in the experimental setup is assumed to be very low.

### Whole genome sequencing and mutation identification

Whole genome sequencing was performed on thirty multi-resistant isolates. Genome sequencing was performed by MicrobesNG (http://www.microbesng.com, Birmingham, UK) according to their protocols (provided in the Supplementary Methods). Trimmed reads were then aligned to a reference genome and variants called using the breseq 0.36.1 pipeline ^112^ also see https://github.com/barricklab/breseq/. The reference genome used was the *E. coli* K-12 BW25113 genome ^98^ NCBI accession CP009273.1, with additional annotations for insertion (IS) element regions to improve the calling of mutations related to IS insertion (modified Genbank format file, Supplementary File 2). One genome sequence did not correspond to *E. coli* K-12 BW25113 and was therefore discarded, leaving *n* = 9 genomes for the ‘low’ mutator treatment and *n* = 10 genomes for the ‘intermediate’ and ‘high’ mutator treatments. Genes relating to drug efflux, drug uptake, and DNA replication and repair were identified using the Ecocyc database https://ecocyc.org, ^113^.

### Fluctuation tests on evolved multi-resistant isolates

Fluctuation tests were performed as in ^109^. Briefly, ancestral BW25113, f..*mutS* and eight evolved multi-resistant strains (four with ≥ 60 mutations identified by whole genome sequencing, and four randomly selected) were streaked from −80°C stocks onto LB agar and incubated overnight at 37°C. One colony each was inoculated into 5 ml of MH broth and grown overnight at 37°C and 250 rpm shaking. Cultures were diluted to an OD of approximately 0.3, and subsequently diluted 1000-fold into MH broth. For each strain, 16 parallel cultures of 300 µl were pipetted into deep-well 96-well plates and grown for 24 h. Each parallel culture was then pipetted onto selective media, i.e. TA agar with 25 µg/l streptomycin. Resistant mutants (*m* counts) were counted after 48 h growth. For each isolate, final population sizes (*N*_*t*_ counts) were estimated from *n* = 3 randomly-selected parallel cultures by serial dilution (to 10^−6^) and plating on TA agar. Mutation rates were estimated from *m* and *N*_*t*_ counts using `mutestim()` from the R package ‘flan’ ^111^.

### Stochastic population dynamics model

We numerically simulated resistance evolution using a stochastic population dynamic model according to the schematic in Figure 3. Model variables and parameters are described in full in Supplementary Table 5 and associated text. Briefly, the model describes four types of resistance *i* ∈ {*S, R, N, D*}, where *S* is antibiotic sensitive, *R* is rifampicin resistant, *N* is nalidixic acid resistant, and *D* is multi-resistant. We write *i*′ to refer to a type *i* with a mutator background (Figure 3). Here we consider a mutator that only differs in mutation rate from the wild-type, hence all growth parameters for types *i* and *i*′ are equal. Simulated populations were initiated with 5.71 × 10^6^ sensitive individuals; this is our estimate of the starting population size in the experiments obtained by serial dilution plating. Of the initial population of sensitive individuals, a fraction *u* were designated mutators, Type *S*′ (*u* = 0, 0.05, 0.1, or 0.3, approximating the observed frequencies in the ‘none’, ‘low’, ‘medium’ and ‘high’ mutator treatments) and the remaining 1 − *u* were designated wild-type, Type *S* (i.e. non-mutators). Bacterial growth was modelled as a Yule-Furry process (see Supplementary Information), but in addition allowing the reproduction rate to depend on population density (which provides equivalent results to continuous-time Gillespie simulations, but with reduced computational time; see Supplementary Figure 8 and associated text.

At the start of the simulation, there were no Type *R*/*R*′, *N*/*N*′, or *D*/*D*′ individuals. Single-resistant Types *R*/*R*′ and *N*/*N*′, individuals must initially arise by mutation in reproduction events of Type *S*/*S*′ individuals. Once single resistant types exist in the population, they may produce further such individuals by reproduction. In reproduction events of individuals of Types *R*/*R*′ or *N*/*N*′, a subsequent mutation can occur, producing Type *D*/*D*′ individuals. Type *D*/*D*′ individuals may also reproduce, but we do not include further mutations in the model. We write *µ*_*R*_/*µ*_*R*_, for the probability with which an offspring acquires resistance to rifampicin by mutation, and *µ*_*N*_ /*µ*_*N*_, for the probability that the offspring acquires resistance to nalidixic acid. We exclude the possibility that both resistance mutations can be newly acquired in the same reproduction event (i.e. no multi-drug resistance mechanisms, and no double-mutation events, which we have shown in the preceding section to be rare).

As the experiment involved a serial transfer protocol, population bottlenecks of 1/200 every 22 h were also simulated. Other parameter values were set to match the experimental procedure (initial frequency of mutators, dilution, duration of experiment; see Supplementary Table 9). Parameter values for growth rates and carrying capacities were estimated from OD growth curves using strains that were derived in an experiment independent of the selection experiments (see Supplementary Figures 4–5) and associated text). The relationship between optical density and colony forming units was used to convert between OD units and numbers of bacteria (see Supplementary Figure 7, Supplementary Table 8 and associated text).

