## Supplementary Information for "Mutators drive evolution of multi-resistance to antibiotics"

### Mutators drive evolution of multi-resistance to antibiotics
### Supplementary Information

Danna R. Gifford, Ernesto Berríos-Caro, Christine Joerres,
Marc Suñé, Jessica H. Forsyth, Anish Bhattacharyya,
Tobias Galla, and Christopher G. Knight

#### Table of contents

**Part 1 Bayesian statistical analysis of experiments and simulations** **S.2**
Detection of resistance during experimental evolution S.2
Growth of strains derived from fluctuation tests S.9
Growth of double resistant strains from selection experiment S.11

**Part 2 Whole genome sequencing and mutation identification** **S.14**
DNA extraction and library preparation S.14
Sequence analysis and mutation calling S.14

**Part 3 Stochastic population dynamics model of resistance evolution** **S.18**
Description of the model S.18
Yule–Furry process and negative binomial distribution S.18
Growth of bacteria S.19
Simulation method S.20
Mutations S.21
Estimating growth parameters from experimental data S.22
Relationship between optical density and bacterial population size S.29
Simulating the experiments S.32
Other simulation conditions S.35

**Part 4 Additional Supplementary Figures** **S.41**

#### Supplementary Information Part 1

### Bayesian statistical analysis of experiments and simulations

#### Detection of resistance during experimental evolution

##### Defining the statistical model

Here we ask whether the initial mutator frequency and antibiotic treatments had an effect on resistance evolution, and whether these effects interacted. We fitted a categorical regression statistical model (also called a 'multinomial logistic' model) to the data, to analyse how different initial mutator frequencies and antibiotic treatments affected which type of resistance was observed at the end of the experiment. Note that in this section, 'model' refers to statistical model, and not to the 'stochastic population dynamics model' introduced later.

To make our analysis explicit, we will briefly describe the statistical model here. We will refer to $Y_i$ as the value of the $i$th observation, $x_{m,i}$ as the independent variables, and $\beta_{m,k}$ as the estimated coefficients. It is the coefficients $\beta_{m,k}$ that are of interest as they relate how different experimental conditions influence the probability of observing any given outcome. Categorical regression can be formulated as an extension of logistic regression. Let $Y_i$ be a categorical variable that takes a value $k$ from $\{1, 2, ..., K\}$, and the probability that $Y_i$ has outcome $k$ be $\mathrm{P}(Y_i = k)$. We use a linear predictor function to compute $\mathrm{P}(Y_i = k)$. For a statistical model considering $M$ explanatory variables, this takes the form

$$f(k, i) = \beta_{0,k} + \beta_{1,k}x_{1,i} + \beta_{2,k}x_{2,i} + ... + \beta_{M,k}x_{M,i}, \quad (1)$$

where $\beta_{m,k}$ is the regression coefficient associated with the $m$th explanatory variable and the $k$th outcome, and $\beta_{0,k}$ is the intercept associated with the $k$th outcome. This function can be written more compactly using vector notation and taking the dot product, $f(k, i) = \vec{\beta}_k \cdot \vec{x}_i$, where $\vec{\beta}_k$ and $\vec{x}_i$ each have length $M + 1$.

The reader may be familiar with binary logistic regression with $K = 2$ outcomes, usually with $k = 1$ defined as 'success' and $k = 0$ as 'failure'. The function $f(k, i)$ is linked to the probability of observing outcome $k$ by taking the log of the odds-ratio of the two outcomes, i.e. the logit function.

If $p_i = \mathrm{P}(Y_i = 1)$,

$$\mathrm{logit}(p_i) = \log\left(\frac{p_i}{1-p_i}\right) = \vec{\beta}_1 \cdot \vec{X}_i$$
$$p_i = \frac{e^{\vec{\beta}_1 \cdot \vec{X}_i}}{1 + e^{\vec{\beta}_1 \cdot \vec{X}_i}}, \qquad (2)$$

where $\vec{X}_i$ is the vector of values taken by the explanatory variables $\vec{x}_i$ for the observation $Y_i$.

For $K > 2$ outcomes, the multinomial logit can be thought of as computing $K-1$ independent logistic regression models with respect to a consistent reference level (Begg and Gray, 1984). If $K$ is chosen as the reference level, $\beta_{0,K}$ is defined as the 'intercept' and all other elements of $\vec{\beta}_K$ are equal to zero. This results in

$$\log\frac{\mathrm{P}(Y_i = 1)}{\mathrm{P}(Y_i = K)} = \vec{\beta}_1 \cdot \vec{X}_i$$
$$\log\frac{\mathrm{P}(Y_i = 2)}{\mathrm{P}(Y_i = K)} = \vec{\beta}_2 \cdot \vec{X}_i$$
$$\cdots \qquad (3)$$
$$\log\frac{\mathrm{P}(Y_i = K-1)}{\mathrm{P}(Y_i = K)} = \vec{\beta}_{K-1} \cdot \vec{X}_i \,.$$

The fact that $\sum_{k=1}^{K} P(Y_i = k) = 1$ allows calculating $\mathrm{P}(Y_i = K) = 1/\left(1 + \sum_{k=1}^{K-1} e^{\vec{\beta}_k \cdot \vec{X}_i}\right)$, which can then be used to solve the other probabilities. For any outcome $c$, the general form of $\mathrm{P}(Y_i = c)$ is thus given as

$$\mathrm{P}(Y_i = c) = \frac{e^{\vec{\beta}_c \cdot \vec{X}_i}}{1 + \sum_{k=1}^{K-1} e^{\vec{\beta}_k \cdot \vec{X}_i}}, \qquad (4)$$

which can then be used to estimate the coefficients $\vec{\beta}_k$ through various methods. The particular method we used, Bayesian categorical regression, is described in the next section.

#### Fitting the statistical model and hypothesis testing using Bayesian categorical regression

In our particular analysis, $Y_i$ represents the type of resistance observed, with five possible outcomes: 'no resistance', 'rifampicin resistance', 'nalidixic acid resistance', 'mixed resistance' and 'double resistance'. For 'mixed resistance', populations grew on selective plates containing either rifampicin or nalidixic acid, but *not* on plates containing both rifampicin and nalidixic acid, whereas 'double resistant' populations grew on plates containing both antibiotics. We note that these outcomes refer to *detection* of resistance, rather than *fixation*, i.e. the frequency of resistant individuals within each population has a value between $> 0$ and $\leq 1$.

The independent variables, $\vec{x}_i$, were the initial mutator frequency, the antibiotic treatment applied, which microtitre plate the population inhabited, and position within each microtitre plate. Initial mutator frequency was treated as a categorical predictor (with levels ‘zero’, ‘low’, ‘medium’ or ‘high’, using ‘zero’ as the reference level), as we have no *a priori* expectation of a linear relationship between the proportion of mutators and the estimated coefficient. Antibiotic treatment was a categorical predictor (with levels ‘no antibiotic’, ‘rifampicin’, ‘nalidixic acid’, or ‘combination’, using ‘no antibiotic’ as the reference level). Initial mutator frequency and antibiotic were each treated as a fixed effect (as variance estimates from random effects variables with fewer than five levels tends to be imprecise, see Harrison, 2015). Plate number and position within each microtitre plate were treated as random effects.

Incorporating both fixed and random effects requires fitting a ‘mixed-effects model’ to the data. However, mixed-effects models for categorical data are not straightforward to fit using standard frequentist inference methods. While it is possible to fit such models in principle (Hedeker, 2003), there is not, to our knowledge, a readily-available software implementation. However, mixed-effects categorical models can be fit using recently-developed tools that use Bayesian inference methods (Bürkner, 2017, 2018). We provide an example below. In addition to the software availability, there are additional benefits of using a Bayesian inference approach (Van Zyl, 2018).

To estimate the coefficients, $\vec{\beta}_k$, we fit a mixed-effects categorical model using `brm()` from the `brms` package (Bürkner, 2017, 2018) in R (R Core Team, 2019). To fit a categorical model, ‘family’ was set to ‘categorical’ (with the default link ‘logit’). To ensure convergence was achieved, we set `max_treedepth` $= 15$ and `adapt_delta` $= 0.99$. We ran four chains of 2000 iterations each, with 1000 burn-in iterations. To permit hypothesis testing on point estimates, samples from specified priors were drawn by setting `sample_prior="yes"`. The choice of priors was based on preliminary data, and is described in detail in a later section. Default values were used for other settings.

To evaluate whether the interaction between initial mutator frequency and antibiotic treatment was important, we compared the full model (i.e. main effects and interaction) to a model with a main-effects only using ‘Pareto-smoothed importance sampling leave-one-out cross-validation’ (PSIS-LOO, Vehtari *et al.*, 2017). Incorporating interactions effects into the model did not significantly improve fit (PSIS-LOO difference in fit: $-2.6 \pm 5.4$ S.E.), hence we present estimates from the main effects model.

An example of the function call is shown below (full R scripts are also available, see ‘Data Availability’ statement in the main text).

```
# Control parameters and priors
priors = c(set_prior ("student_t(7, -5, 2.5)", class = "Intercept"),
      set_prior ("student_t(7, 0, 2.5)", class = "b"))
controls = list(adapt_delta = 0.99, max_treedepth = 15)
# Model calls
M1.full = state ~ (pmutS.text + antibiotic)^2 + (1|row) + (1|col)
M1.main = state ~ (pmutS.text + antibiotic)   + (1|row) + (1|col)
```

```
        modelM1.full = brm(M1.full,
              family = categorical("logit"),
              chains = 4, cores = 4, iter = 2000, warmup = 1000,
              prior = priors, control = controls, sample_prior = "yes",
              data = popnsday6)

        modelM1.main = brm(M1.main,
              family = categorical("logit"),
              chains = 4, cores = 4, iter = 2000, warmup = 1000,
              prior = priors, control = controls, sample_prior = "yes",
              data = popnsday6)
```

#### Establishing priors for the model

Priors were established through a preliminary experiment in which pure populations of either wild-type or mutator bacteria were subjected to increasing concentrations of single and combination antibiotic treatments (see Methods in the main text). For this preliminary experiment, we assayed the proportion of populations by measuring OD at 600 nm relative to a 'blank' well with no bacteria present, using a BMG POLARstar OPTIMA plate reader (BMG Labtech, Ortenberg, Germany). A population was considered to be 'alive' with OD $> 0.1$, otherwise 'extinct'. The data are presented in Supplementary Figure 1. We observed that, in the absence antibiotic treatment, all populations survived. For single antibiotic treatments, all mutator populations survived, but some wild-type populations went extinct. For the combination antibiotic treatment, 0/54 wild-type and 39/54 mutator populations survived. We use this information to first establish priors for the intercepts of the model (Supplementary Figure 1B).

As we observed no alive wild-type populations in the combination treatment, we therefore have some confidence that the proportion of multi-resistant outcomes is then $p_{\text{double}} < 1/54$. Therefore, the intercept for multi-resistance should be less than $\text{logit}(1/54) \approx -3.97$. We therefore used a $t$-distribution with mean $\mu = -5$ and broad and heavy tails with standard deviation $\sigma = 2.5$, and degrees of freedom $\nu = 7$, to reflect uncertainty. This distribution covers both $p_{\text{double}} \approx 0$ in the left tail, and $p_{\text{double}} = 39/54$, i.e. the number of alive populations in pure mutator populations, in the right tail. Given that the probability of being single-drug resistant in the absence of antibiotic is likely greater than double resistant, intercepts for other resistance outcomes are likely covered by this distribution, and so we use the same prior for the single-resistance intercepts.

Next, we use this information to establish priors for the other coefficients. We observed that the presence of mutators increases survival in the presence of both single and combination antibiotics, hence the effect of mutators on *total* resistance (i.e. the sum of all resistance states observed) is likely to be positive. However, it is possible that different resistance states may have a negative relationship with the proportion of mutators, if for example, elevated mutation rates pushes double resistance to arise in the background of single-drug resistance, single-drug

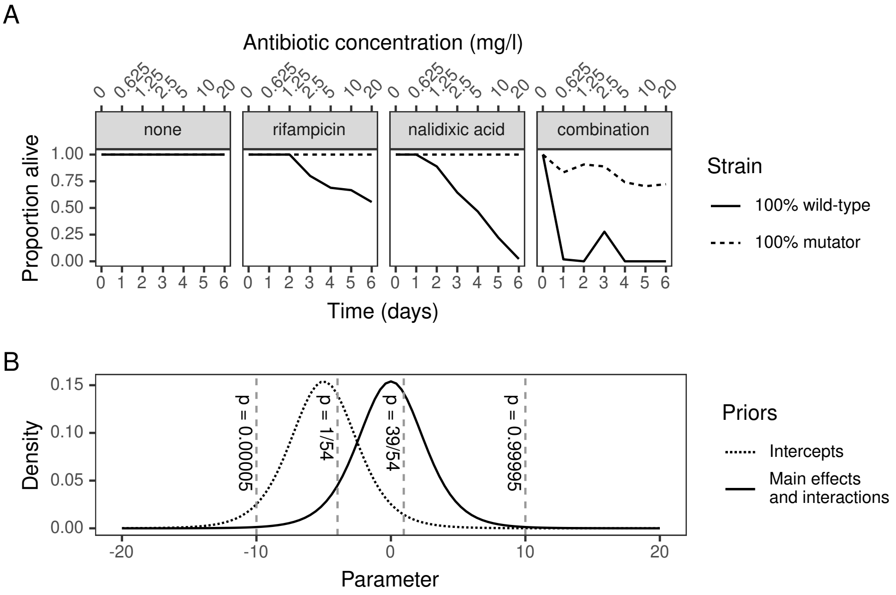


**Supplementary Figure 1:** A. Preliminary experiment tracking the proportion of populations 'alive' (i.e. with OD $> 0.1$) under experimental treatments. B. Illustration of priors used to conduct Bayesian categorical regression using information from preliminary experiment, indicated by vertical dashed lines. Curves show the probability density for a Student's $t$ distribution with means $\mu = 0$ (solid) or $\mu = -5$ (dotted), standard deviation $\sigma = 2.5$ (both curves) and degrees of freedom $\nu = 7$ (both curves).

resistance may have a negative relationship with initial mutator frequency. Hence for the effects of initial proportion of mutators, we use a $t$-distribution centred at $\mu = 0$ with the same broad and heavy tails $\sigma = 2.5$, and $\nu = 7$ to reflect uncertainty. This covers the observed proportion of alive populations in a purely mutator population $p = 39/54$; we should however expect to observe fewer resistance events when the proportion of mutators is less than 1, as was the case in the selection experiment described in the main text. It also allows for the extreme possibilities of (nearly) zero or (nearly) all resistance, albeit with less weight given.

From this experiment, we have limited direct evidence for how the presence of antibiotics should affect the probability of observing resistance. On one hand, the presence of antibiotics decreased the number of alive populations (and an extinct population cannot be resistant). On the other, our growth measurements of resistant strains in the presence of antibiotics suggest a positive effect of being resistant in the presence of antibiotics, which would allow them to spread to high frequency and thus escape loss due to genetic drift. The effect of antibiotics is likely to be in the same range as for mutators (which includes 'no effect'), hence we use the same prior distribution (i.e. a $t$-distribution with $\mu = 0$, $\sigma = 2.5$, and $\nu = 7$).

#### Estimated model coefficients

Estimated model coefficients for the main-effects only model are shown in Supplementary Table 1. These are reported as treatment contrasts (i.e. relative to the treatment of no antibiotics and no mutators). We used 95% credible intervals (95% C.I.s) for all hypotheses tested. Coefficients were estimated on the logit scale (i.e. log-odds, which can assume any value between $-\infty$ and $\infty$, corresponding to proportions of outcomes of $p_{\text{resistance state}} = 0$ and $1$ respectively). A coefficient is estimated for each possible combination of response outcome (resistance state) and the predictors (initial mutator frequency and antibiotic), which indicates how a specific combination of predictor levels (e.g. 'low' mutator frequency and 'rifampicin' treatment) influences the probability that a given resistance state is observed (e.g. 'rifampicin resistance').

Mutators generally have a positive effect on resistance, as indicated by positive treatment contrasts (Supplementary Table 1). However, as previously noted, although we expect an overall positive association between mutator frequency and resistance, we do not necessarily expect a straightforward relationship with each resistance state. This is because the 'mixed resistance' and 'double resistance' states follow from a single-drug resistance state. Indeed, this is the case for the 'low' mutator treatment and nalidixic acid resistance state, which is overtaken by 'mixed resistance'. Recall that the mixed resistance state contains both single-drug resistance types. Thus, to assess the effect of mutators on the presence of nalidixic acid resistance types, we can combine these treatment contrasts. We can perform Bayesian hypothesis tests on combinations of treatment contrasts using `hypothesis()`. The combined treatment contrast for the 'low' mutator frequency for the combination of 'nalidixic acid resistance' and 'mixed resistance' is 3.36 [95% C.I. of (1.74, 5.00)], which is positive as predicted. The same combination of treatment contrasts could be performed for rifampicin resistance or for the other initial mutator frequencies, but as these already exclude zero, the combinations of their contrasts will also exclude zero.

**Supplementary Table 1:** Effects of initial mutator frequency and antibiotic treatment on resistance state observed at the end of the experiment. Estimated model coefficients come from fitting a Bayesian categorical regression model to the selection experiment outcomes. Treatment contrasts on the logit scale are shown. (* denotes 95% credible intervals excluding zero).

| Resistance state | Coefficient | Treatment contrast | 95% credible interval | |
|---|---|---|---|---|
| rifampicin resistance | intercept | -2.83 | (-3.58, -2.14) | * |
| | low | 1.48 | (0.78, 2.25) | * |
| | intermediate | 1.98 | (1.23, 2.74) | * |
| | high | 3.36 | (2.56, 4.17) | * |
| | rifampicin | 3.29 | (2.60, 4.02) | * |
| | nalidixic acid | -1.22 | (-2.28, -0.29) | * |
| | combination | -2.22 | (-3.16, -1.38) | * |
| nalidixic acid resistance | intercept | -4.00 | (-5.06, -3.08) | * |
| | low | -0.88 | (-2.30, 0.34) | |
| | intermediate | 1.51 | (0.65, 2.39) | * |
| | high | 2.81 | (1.90, 3.78) | * |
| | rifampicin | -2.41 | (-7.70, 0.75) | |
| | nalidixic acid | 2.67 | (1.88, 3.54) | * |
| | combination | -0.88 | (-2.11, 0.26) | |
| mixed resistance | intercept | -6.78 | (-8.46, -5.37) | * |
| | low | 4.27 | (3.02, 5.81) | * |
| | intermediate | 4.89 | (3.60, 6.45) | * |
| | high | 6.51 | (5.20, 8.10) | * |
| | rifampicin | 3.63 | (2.77, 4.52) | * |
| | nalidixic acid | 2.84 | (2.18, 3.53) | * |
| | combination | -0.96 | (-1.90, -0.10) | * |
| double resistance | intercept | -10.99 | (-13.91, -8.71) | * |
| | low | 5.14 | (3.57, 7.39) | * |
| | intermediate | 5.98 | (4.36, 8.26) | * |
| | high | 8.01 | (6.35, 10.28) | * |
| | rifampicin | 7.80 | (6.21, 9.97) | * |
| | nalidixic acid | 5.20 | (3.71, 7.36) | * |
| | combination | 4.22 | (2.78, 6.33) | * |

#### Checking robustness against choice of priors

A potential consequence of using informative priors in Bayesian inference is that they may influence the posterior distribution for estimated coefficients unduly if they are not chosen appropriately. However, the use of non-informative priors does not necessarily mitigate these problems (for an in-depth discussion, see Lemoine, 2019). To check the robustness of our model against the originally chosen priors, we refit the model using different priors. We set Student-$t$ priors on the intercept with arbitrarily chosen $\mu$ of -10, -20, -30, -40, with (as before) $\sigma = 2.5$ and $\nu = 7$. The estimated coefficients resulting from using these priors are quantitatively similar to our original model.

As a second approach to evaluating robustness, we also set strongly-informative priors on each coefficient of the model. We estimated means for each coefficient using a fixed-effects categorical model (i.e. without the random effects) by maximum likelihood using the `multinom()` function from the nnet package (Venables and Ripley, 2002), which refers to this type of model as 'multinomial logistic regression'. Note that assigning priors in this fashion is used here only as a diagnostic technique, and is not recommended as a basis for assigning priors more generally. For the model with these strong priors, the majority of coefficients were again similar to the model with weaker priors. The exception was for coefficients associated with the 'double resistance' outcome. Here, because there were zero double resistance events associated with the reference levels of the main effects of the model (i.e. no mutators, no antibiotics), the maximum likelihood estimated the intercept associated with this outcome to be very small (log-odds of $-38.34$, equivalent to an odds ratio of approximately $4.5 \times 10^{-16}$). The posterior distribution for the intercept was dominated by this strong prior. Consequently, the estimated coefficients for the coefficients associated with antibiotic treatment and the presence of mutators where there were double resistant outcomes observed were much larger than those estimated when using the original weakly-informative priors. However, we note that in all cases, the qualitative outcomes with respect to antibiotic treatment and the presence of mutators remain unchanged.

### Growth of strains derived from fluctuation tests

#### Defining the statistical model

Here we determined under which conditions resistant strains achieved a growth advantage (see Supplementary Figure 3B). Recall that growth was characterised using area under the curve (AUC) of growth curves from OD measured at 600 nm. We compare the wild-type (*E. coli* K-12 BW25113) with single- and double-drug resistant mutants selected in the BW25113 genetic background through fluctuation tests (see Methods in main text). The model fitted to the data in Figure 2 is a Bayesian two-way factorial model, with 'strain' and 'antibiotic' as predictor variables. We treated AUC measured in each antibiotic treatment as a multivariate response. This was on the basis that each strain was measured at several different concentrations, hence may be non-independent. We used Student's $t$ priors, as opposed to a Gaussian priors, because doing so is more robust against extreme values (i.e. values that appear to be 'outliers', but where there is no evidence of error in data collection, FENG *et al.*, 2015).

#### Establishing priors for the statistical model

The intercept of this model is the mean AUC from the growth curves of OD, which is always greater than zero. To calculate empirical AUC, we used `SummariseGrowth()` from the `growthcurver`, which uses the trapezoidal rule to approximate the integral under the curve. OD values on this BMG FLUOstar OPTIMA plate reader are typically less than 1.2 for blank-corrected values. We used the trapezoidal rule to calculate extremes for the values of AUC, i.e.

$$\text{AUC} = \int_a^b f(x)dx \approx (b-a)\frac{f(a)+f(b)}{2}.$$

Since AUC was calculated over 25 h, the lower extreme is $(25-0)(0+0)/2 = 0$ (i.e. no growth) and the upper extreme is $(25-0)(1.2+1.2)/2 = 30$ (i.e. essentially instantaneous achievement of maximum density). However, previous experience from growing wild-type *E. coli* suggests they exit exponential growth in the region of 10 h after inoculation after a 1/1000 dilution in MH broth, giving

$$\text{AUC} \approx \frac{(10-0)(0+1.2)}{2} + \frac{(25-10)(1.2+1.2)}{2} = 24$$

as a rough approximation of what would be expected under good growth conditions. Hence, the prior for the intercepts should give highest density between 0 and 24. This was specified as a $t$-distribution with mean $\mu = 10$ and broad and heavy tails with standard deviation $\sigma = 2.5$, and degrees of freedom $\nu = 7$. The presence of mutators could either increase or decrease evolved fitness relative to the intercept. To set the priors on the effect of mutators, we use a $t$-distribution centred on mean $\mu = 0$, with standard deviation $\sigma = 0.5$, and degrees of freedom $\nu = 7$. If the intercept takes a value in the region of the mean of its prior, this prior on the mutator effect allows for the extreme possibility that the presence of mutators results in an AUC of zero (if the coefficient takes value $-10$), or an AUC beyond the technical capabilities of the equipment (for values $> 10$), with low probability. This is illustrated in Supplementary Figure 2.

#### Fitting the statistical model and hypothesis testing

As previously, model fitting was performed using `brm()`. To use a Student's $t$ model, 'family' was set to 'student' (with the default link 'identity'). To permit hypothesis testing on point estimates, samples from specified priors were drawn by setting `sample_prior="yes"`. To ensure convergence, we set `max_treedepth` $= 15$ and `adapt_delta` $= 0.99$. Default values were used for other settings. As before, we used a 95% C.I. for hypotheses. As we are primarily interested in hypotheses on point estimates, we do not present all of the estimated effects here, though this can be generated with the R script provided (see the 'Data Availability' statement in

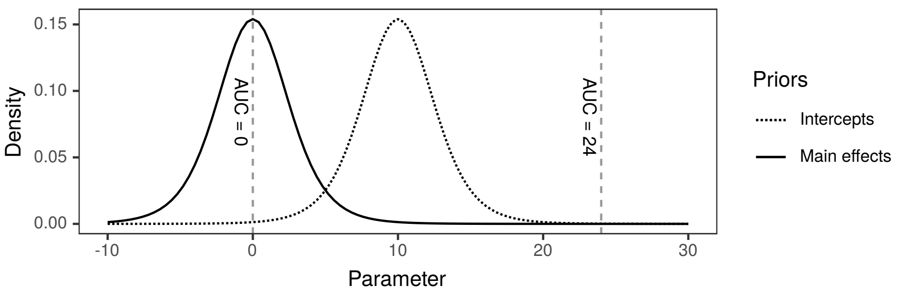


**Supplementary Figure 2:** Illustration of priors used to conduct Bayesian multivariate regression on the effect of mutators on growth. Curves show the probability density for a Student's $t$ distribution with means $\mu = 0$ (solid) or $\mu = 10$ (dotted), standard deviation $\sigma = 2.5$ (both curves) and degrees of freedom $\nu = 7$ (both curves). Dashed vertical lines show prior information on technical upper and lower values for AUC, used to set a prior on the intercept.

the main text). We calculated the difference in AUC of the single-resistant and double-resistant strains in the two single-drug treatments. Estimated model coefficients are reported as treatment contrasts (i.e. relative to the wildtype and antibiotic-free treatment). For the rifampicin treatment, there was no benefit of double resistance over rifampicin resistance [95% C.I. of the difference = $-0.07$, 95% C.I.: $(-0.38, 0.23)$]. For the nalidixic acid treatment, there was a deleterious effect of double resistance over nalidixic acid resistance [95% C.I. of the difference = $-0.75$, 95% C.I.: $(-1.05, -0.45)$].

### Growth of double resistant strains from selection experiment

#### Defining the statistical model

Here we determine whether initial mutator frequency had an effect on the growth of double resistant strains that evolved during selection. Decreased growth with a higher mutator frequency may occur if deleterious variation accumulated by the elevated mutation rate hitch-hikes to high frequency along with resistance. Alternatively, increased growth may be realised if the elevated mutation rate allowed the accumulation of beneficial mutations, or increased the clonal interference among resistance mutations. We fit a Bayesian multivariate Student's $t$ mixed-effects model. We treated growth (measured by AUC) in the presence (20 mg/l) and absence (0 mg/l) as a bivariate response variable, and initial mutator frequency as the sole population-level factor ('low', 'intermediate', 'high', with the growth of double resistant strains derived in the BW25113 wild-type background as the reference level). As before, we used a bivariate model to account for correlations arising from measuring the AUC of each strain multiple times in different environments and a Student's $t$ model to be robust to 'outliers' in the data. AUC at each concentration was measured over several 'replicate' experiments, which was treated as a varying factor common to both response variables.

#### Establishing priors for the bivariate linear model

The same experimental procedure was used to measure growth of the selection experiment strains, hence we use the same priors as for the fluctuation test-derived strains (Supplementary Figure 2).

#### Fitting the statistical model and hypothesis testing

As previously, statistical model fitting was performed using `brm()`. To use a Student's $t$ model, 'family' was set to 'student' (with the default link 'identity'). To permit hypothesis testing on point estimates, samples from specified priors were drawn by setting `sample_prior="yes"`. To ensure convergence, we set `max_treedepth` $= 15$ and `adapt_delta` $= 0.99$. Default values were used for other settings. As before, we used 95% C.I.s for hypotheses. Estimated coefficients are given in Supplementary Table 2, reported as treatment contrasts (i.e. relative to double resistance in the BW25113 strain). Growth measured by AUC in 0 mg/l and 20 mg/l of the combination antibiotic treatment was positively correlated [$r = 0.68$, 95% C.I.: $(0.56, 0.79)$]. The statistical model incorporating initial mutator frequency was a worse fit than an intercept-only model (WAIC $564.6 \pm 23.7$ SE vs. $540.0 \pm 25.2$ SE), suggesting initial mutator frequency did not have a large influence on AUC of double resistant strains.

**Supplementary Table 2:** Estimated coefficients for the population-level effect of mutators on growth (measured by AUC) of multi-resistant clones in 0 mg/l and 20 mg/l of the combination treatment, from the fit of a Bayesian bivariate regression model. Treatment contrasts are shown (* denotes 95% credible intervals excluding zero).

| Concentration | Coefficient | Estimate | Error | 95% credible interval | |
|---|---|---|---|---|---|
| AUC in 0 mg/l | intercept | 8.70 | 0.47 | $(7.71, 9.58)$ | * |
| | low | -1.68 | 0.59 | $(-2.74, -0.43)$ | * |
| | intermediate | -0.51 | 0.56 | $(-1.55, 0.64)$ | |
| | high | -0.58 | 0.53 | $(-1.54, 0.53)$ | |
| | $\sigma$(intercept) | -1.75 | 0.45 | $(-2.60, -0.83)$ | * |
| | $\sigma$(low) | 0.57 | 0.50 | $(-0.44, 1.53)$ | |
| | $\sigma$(intermediate) | 0.07 | 0.50 | $(-0.95, 1.05)$ | |
| | $\sigma$(high) | 0.14 | 0.46 | $(-0.82, 1.02)$ | |
| AUC in 20 mg/l | intercept | 7.20 | 0.46 | $(6.27, 8.05)$ | * |
| | low | -0.77 | 0.57 | $(-1.85, 0.39)$ | |
| | intermediate | 0.17 | 0.55 | $(-0.93, 1.26)$ | |
| | high | -0.01 | 0.51 | $(-0.94, 1.02)$ | |
| | $\sigma$(intercept) | -2.13 | 0.48 | $(-3.07, -1.12)$ | * |
| | $\sigma$(low) | 0.46 | 0.52 | $(-0.61, 1.44)$ | |
| | $\sigma$(intermediate) | 0.41 | 0.53 | $(-0.69, 1.43)$ | |
| | $\sigma$(high) | 0.39 | 0.48 | $(-0.64, 1.30)$ | |

#### Supplementary Information Part 2

### Whole genome sequencing and mutation identification

#### DNA extraction and library preparation

We performed whole genome sequencing on isolates from thirty of the evolved multi-resistant mutator populations. Genome sequencing was performed by MicrobesNG (`http://www.microbesng.com`, Birmingham, UK) according to the following protocols. Pure cultures of each strain were grown as a lawn covering approximately 1/3 of a 90mm Petri dish, with additional streaking on the remaining 2/3 to ensure culture purity. Cells were harvested and resuspended in DNA/RNA Shield (Zymo Research, USA). From each suspension, 5 to 40 µl was lysed with 120 µl of TE buffer containing lysozyme (final concentration 0.1 mg/mL) and RNase A (ITW Reagents, Barcelona, Spain) (final concentration 0.1 mg/mL), incubated for 25 min at 37°C. Proteinase K (VWR Chemicals, Ohio, USA; final concentration 0.1mg/mL). SDS (Sigma-Aldrich, Missouri, USA) (final concentration 0.5% v/v) was added and incubated for 5 min at 65°C. Genomic DNA was purified using an equal volume of SPRI beads (Beckman Coulter, USA) and resuspended in EB buffer (Qiagen, Germany). DNA is quantified with the Quant-iT dsDNA HS kit (ThermoFisher Scientific) assay in an Eppendorf AF2200 plate reader (Eppendorf UK Ltd, United Kingdom). Genomic DNA libraries were prepared using the Nextera XT Library Prep Kit (Illumina, San Diego, USA) following the manufacturer's protocol with the following modifications: input DNA is increased 2-fold, and PCR elongation time is increased to 45 s. DNA quantification and library preparation are carried out on a Hamilton Microlab STAR automated liquid handling system (Hamilton Bonaduz AG, Switzerland). Pooled libraries are quantified using the Kapa Biosystems Library Quantification Kit for Illumina. Libraries are sequenced using Illumina sequencers (HiSeq/NovaSeq) using a 250bp paired end protocol. Adapters were trimmed from reads using Trimmomatic 0.30 with a sliding window quality cutoff of Q15 (Bolger *et al.*, 2014).

#### Sequence analysis and mutation calling

Trimmed reads were then aligned to a reference genome and variants called using the breseq 0.36.1 pipeline (`https://github.com/barricklab/breseq/`, Deatherage and Barrick, 2014). The reference genome used was the *E. coli* K-12 BW25113 genome (Grenier *et al.*, 2014), with additional annotations for insertion (IS) element regions to improve the calling of mutations related to IS movement (modified Genbank available in the supplementary information). One isolate was discarded after sequencing due to being the wrong organism, ultimately leaving for the analysis $n = 9$ genomes for the 'low' mutator treatment and $n = 10$

genomes for the ‘intermediate’ and ‘high’ mutator treatments. Mutations from all isolates were brought together into a single file using the breseq-provided utility program `gdtools ANNOTATE`. Mutations in canonical targets of resistance and efflux pumps, as well as the total number of mutations detected in each isolate, are given in Supplementary Table 3. Separate output files generated by breseq are provided as Supplementary File 1. Mutations relating to efflux pumps were identified using the Ecocyc database (`https://ecocyc.org`, KESELER *et al.*, 2011).

We performed whole genome sequencing and variant calling on multi-resistant isolates. All 29 isolates acquired rifampicin resistance mutations in the canonical mutational target *rpoB*; for nalidixic acid resistance, 26/29 acquired mutations in the canonical target *gyrA*, and a further 3/29 in *gyrB*. Diverse mutations were observed in both of these targets (Supplementary Table 3), with the most prevalent being *rpoB* D516G and *gyrA* D72G. There was no particular association between specific SNP occurring in *rpoB* and *gyrA*, excepting the pair *rpoB* D516G and *gyrA* D72G, which occurred 5 times in total. In addition to mutations in these canonical targets, most strains (21/29) acquired a mutation in one of the multi-drug efflux pump systems in *E. coli*, the most frequently hit being *acrR* (9/29), a repressor involved in the AcrAB-TolC system. However, whether all such mutations are functionally significant is unclear, as these included both synonymous substitutions and putative loss-of-function mutations in structural components, which are not likely to improve efflux. In addition to resistance-associated mutations, we detected additional mutations across the genome (median = 34, range = 25–162, Figure 2C), with a mutational spectrum of SNPs consistent with $\Delta$*mutS* (Figure 2D). While some likely affected fitness, it is likely that many mutations observed were selectively neutral, as 283/1252 (22.6%) of all mutations detected were synonymous SNPs. Notably, in 4/29 isolates, we identified a greater than average number of mutations (from the ‘low‘ treatment: three isolates with 62, 99, and 162 mutations each, and from the ‘intermediate‘ treatment: one isolate with 60 mutations). Among these, we detected mutations in DNA replication genes (Supplementary Table 4), indicating a potential advantage to an even higher mutation rate for acquiring multi-resistance.

**Supplementary Table 3:** Resistance-associated mutations detected in multi-resistant isolates that evolved via ramping selection in the combination treatment.

| Mutator frequency | ID | Rifampicin resistance | Nalidixic acid resistance | Efflux mutations | Total mutations |
|---|---|---|---|---|---|
| low | B7 | *rpoB* D516G | *gyrA* D72G | *acrR* A20V | 39 |
| | C6 | *rpoB* S512P | *gyrA* D87A | *macB* C580Y, *mprA* R111C | 31 |
| | C10 | *rpoB* Q148R, *rpoC* G336S | *gyrB* D426G | *acrB* G675G, *mprA* 125fs | 62 |
| | E10 | *rpoB* P564L | *gyrA* D87G | *acrB* 23fs, *mdtK* W136R, *mdtE* R65R | 162 |
| | E4 | *rpoB* L511P | *gyrA* D87G | *acrA* Q36X, *acrR* T44A, *emrK* D41G, *marR* G116D | 33 |
| | F4 | *rpoB* S512P | *gyrA* D82G | between *acrA*/acrR [intergenic (-95/-47)] | 26 |
| | F6 | *rpoB* G534S | *gyrA* E153G | *acrR* P206L, *marR* L97P, *mdtO* R542R | 45 |
| | G3 | *rpoB* I572F | *gyrA* G75S | *acrA* V371A, *mprA* G121D | 99 |
| | G9 | *rpoB* L511P | *gyrA* D72G, *gyrB* Y483C | *marR* A70T | 34 |
| intermediate | B4 | *rpoB* L511P | *gyrA* D72G | *emrK* D41G, *marR* A70T, *mprA* L29P, *tolC* L234L, *rob* A70V | 39 |
| | B5 | *rpoB* G534S | *gyrA* G81D | *emrK* D41G, *mprA* S84P, *mdtL* 186fs | 60 |
| | B6 | *rpoB* G534D | *gyrA* A51V | *mprA* R111H | 45 |
| | C4 | *rpoB* D516G | *gyrA* D72G | *acrR* 29fs, *marR* 126fs, *rpoS* K204E | 29 |
| | C10 | *rpoB* D516G | *gyrA* D72G | *acrR* 189fs, *marR* 62fs | 36 |
| | D10 | *rpoB* Q148R | *gyrA* S83L, *gyrB* P747S | none | 32 |
| | E2 | *rpoB* L511P | *gyrA* S83L | *acrR* T44A | 25 |
| | E4 | *rpoB* S512P | *gyrA* S83L | none | 28 |
| | G2 | *rpoB* D516G | *gyrA* D72G | *acrR* R13H | 37 |
| | G9 | *rpoB* S512P | *gyrA* D87G, *gyrB* R291R | none | 30 |
| high | C4 | *rpoB* Q148R | *gyrA* D87G | *marR* R77H | 40 |
| | C11 | *rpoB* G534D | *gyrB* D426G | *acrR* A41V, *emrK* D41G | 32 |
| | D8 | *rpoB* D516G | *gyrA* S83L | *acrR* S172P | 26 |
| | E8 | *rpoB* G534S | *gyrA* D72G | *marR* 62fs, *mprA* 97fs | 41 |
| | F3 | *rpoB* Q148R | *gyrA* D72G | *acrR* 29fs | 48 |
| | F5 | *rpoB* S512P | *gyrA* S83P | none | 34 |
| | F10 | *rpoB* L533P | *gyrA* D72G | *mprA* L41P | 44 |
| | G5 | *rpoB* P564L | *gyrA* D87N | none | 31 |
| | G7 | *rpoB* S512P | *gyrB* K447E | none | 31 |
| | G11 | *rpoB* D516G | *gyrA* D72G | *acrR* 189fs | 33 |

X–mutation to premature stop codon; fs–frameshift mutation

**Supplementary Table 4:** Increases in mutation rate associated with mutations in known DNA replication and repair genes.

| Mutator frequency | ID | Gene | Product | Mutation type | Total mutations | Fold change in $\mu$ |
|---|---|---|---|---|---|---|
| low | E10 | *dnaK* | chaperone Hsp70, with co-chaperone DnaJ | S | 162 | 51.1 |
| | | *dnaQ* | DNA polymerase III epsilon subunit | NS | | |
| | | *umuD* | DNA polymerase V, subunit D | S | | |
| | | *recN* | recombination and repair protein | NS | | |
| | | *sbmC* | DNA gyrase inhibitor | stop | | |
| | | *ligA* | DNA ligase, NAD(+)-dependent | S | | |
| low | C6 | — | — | — | 31 | 11.5 |
| low | E4 | *polB* | DNA polymerase II | NS | 33 | 9.07 |
| low | G3 | *nei* | endonuclease VIII | NS | 99 | 3.57 |
| | | *lon* | DNA-binding ATP-dependent protease La | indel | | |
| | | *mcrA* | type IV methyl-directed restriction enzyme (e14 prophage) | NS | | |
| | | *ung* | uracil-DNA-glycosylase | S | | |
| | | *smf* | DNA recombination-mediator A family protein | NS | | |
| low | C10 | *dnaJ* | chaperone Hsp40, DnaK co-chaperone | NS | 62 | 3.03 |
| | | *polB* | DNA polymerase II | indel | | |
| low | G9 | *cho* | endonuclease of nucleotide excision repair | NS | 34 | 0.956 |
| intermediate | E2 | *nfo* | endonuclease IV with intrinsic 3'-5' exonuclease activity | NS | 25 | 0.932 |
| intermediate | B5 | *uvrB* | excinulease of nucleotide excision repair | NS | 60 | 0.467 |

NS–non-synonymous SNP; S–synonymous SNP; stop–premature stop codon; indel–insertion/deletion mutation

#### Supplementary Information Part 3

### Stochastic population dynamics model of resistance evolution

#### Description of the model

We numerically simulated resistance evolution using a stochastic population dynamic model. A conceptual diagram of the model is shown in Figure 3 of the main text. The model describes four strains $i \in \{S, R, N, D\}$, where $S$ is the sensitive ancestor, $R$ is rifampicin resistant, $N$ is nalidixic acid resistant, and $D$ is double resistant.

Populations initially consist only of type $S$, a fraction $u$ of which are mutators and $1-u$ of which are wild-type. The conditions of the simulation are intended to replicate the selection experiment (see main text Methods for details). The simulated populations experience increasing antibiotic concentrations, which double each day. As in the experiment, populations are allowed to grow for a fixed time period, during which mutations can arise, before being subjected to dilution. These processes are described in the following sections.

#### Yule–Furry process and negative binomial distribution

To simulate the growth of bacteria, we assume that each of the $n_i$ individuals of strain $i$ replicates at any time at a certain rate, regardless of its age. This rate will depend on the strain $i$, and on the composition of the population, as described below. For the time being we simply write $\lambda_i$ for this rate. Setting mutations aside for the moment and assuming that replication events are independent, this is described by the well-known Yule-Furry process. The corresponding master equation reads

$$\dot{p}_{n_i}(t) = -n_i \lambda_i p_{n_i}(t) + (n_i - 1)\lambda_i p_{n_i - 1}(t), \tag{5}$$

where $p_{n_i}(t)$ is the probability that the number of bacteria of type $i$ is $n_i$ at time $t$. Here $\dot{p}_{n_i}$ represents the first derivative of the probability with respect to time. The first term on the right hand side of Eq. (5) accounts for the outflux of probability (per unit time) of state $n_i$, and the second for the influx into state $n_i$. We denote the solutions of Eq. (5) by $p_{n_i}(t; n_i^0)$ where $n_i^0$ indicates the number of bacteria at $t = 0$.

When the rate $\lambda_i$ is constant in time the master equation Eq. (5) can be solved analytically. The solution with initial condition $p_{n_i}(0; n_i^0) = \delta_{n_i, n_i^0}$ (where $\delta_{n_i, n_i^0}$ is the Kronecker delta) is the negative binomial distribution

$$p_{n_i}(t; n_i^0) = \binom{n_i - 1}{n_i - n_i^0} e^{-\lambda_i n_i^0 t} \left(1 - e^{-\lambda_i t}\right)^{(n_i - n_i^0)} = \binom{n_i - 1}{n_i - n_i^0} p_i^{n_i^0} q_i^{(n_i - n_i^0)}, \tag{6}$$

where $p_i = e^{-\lambda_i t}$and $q_i = 1 - p_i$. The expression in Eq. (6) is the probability that there are $n_i$ individuals of strain $i$ in the population at time $t$, given that there were $n_i^0$ such individuals at time $t = 0$. This does not involve any approximation for a pure birth process with constant reproduction rate $\lambda_i$.

### Growth of bacteria

#### Saturated growth

In the following we write $n_T = n_S + n_R + n_N + n_D$ for the total number of bacteria in the population. In order to capture the saturation of growth found in the experiment we use the following expression for the reproduction rates $\lambda_i$ for strain $i$ when $n_T \leq k_i$:

$$\lambda_i = r_i \left(1 - \frac{n_T}{k_i}\right). \tag{7}$$

The parameter $k_i$ is the carrying capacity for strain $i$, and represents the maximum value $n_i$ can take in absence of any other strain. The pre-factor $r_i$ is the growth rate per unit time of strain $i$ when there are very few individuals in the population (i.e., when the suppression of growth has not yet set in). The expression $r_i(1 - n_T/k_i)$ accounts for the interaction between the different strains. Terms of this type are common in Lotka–Volterra type models. In writing down the above expression for $\lambda_i$ we have assumed that the interaction is the same between each pair of strains (pure scramble competition, i.e. no direct interference, cross-feeding or similar). Given that $n_T$ is time dependent, the reproduction rate $\lambda_i$ is time dependent as well.

We interpret the reproduction rate $\lambda_i$ as an effective quantity, taking into account both birth and death processes of bacteria. We set the reproduction rate $\lambda_i$ to zero when $n_T$ exceeds $k_i$. This guarantees that the growth of strain $i$ saturates when $n_T = k_i$, and at the same time it prevents the species with the highest carrying capacity from automatically taking over the population, as explained in more detail in (Berríos-Caro *et al.*, 2021).

#### Connection to Lotka–Volterra dynamics

To make the connection to the well-known Lotka-Volterra dynamics more explicit, we note that the expected value of $n_i$ obtained from the distribution in Eq. (6) is $\bar{n}_i(t) = n_i^0 e^{\lambda_i t}$. This means that the mean number of individuals $\bar{n}_i(t)$ at time $t$ fulfills the relation $\mathrm{d}\bar{n}_i/\mathrm{d}t = \lambda_i \bar{n}_i$. With the choice of $\lambda_i$ as in Eq. (7) this turns into a competitive Lotka-Volterra equation

$$\frac{\mathrm{d}\bar{n}_i}{\mathrm{d}t} = r_i \bar{n}_i \left(1 - \frac{\bar{n}_T}{k_i}\right). \tag{8}$$

**Supplementary Table 5:** Definitions of model variables and parameters.

| Parameter | Definition |
|---|---|
| $t$ | time (hours) |
| $i$ | strain $i \in \{S, R, N, D\}$ |
| $n_i$ | number of individuals (of strain $i$) |
| $\bar{n}_i$ | mean number of $n_i$ over different realisations of the experiment |
| $r_i$ | growth rate of strain $i$ |
| $k_i$ | carrying capacity of strain $i$ |
| $\mu_j$ | mutation rate to resist antibiotic, $j \in \{R, N\}$ |
| $n_T$ | total population size, $n_T = \sum_i n_i$ |
| $\bar{n}_T$ | mean of $n_T$ over different realisations of the experiment |
| $\Delta t$ | length of time per simulation time-step |
| $n^c$ | critical population size at growth regime switch |
| $t_c$ | time at growth regime switch |
| $u$ | initial frequency of mutators |

If we consider a a 'pure culture' where only a single type of bacterium is present, i.e. $\bar{n}_i = \bar{n}_T$, this has the analytical solution

$$\bar{n}_i(t) = \frac{k_i \bar{n}_i(t_0)}{\bar{n}_i(t_0) + [k_i - \bar{n}_i(t_0)]e^{-r_i(t-t_0)}} \quad . \tag{9}$$

Eq. (9) describes a sigmoidal dynamic, starting from $\bar{n}_i(t_0)$ at time $t_0$, and approaching an asymptotic limit at the carrying capacity $k_i$ at long times. This is the familiar logistic growth model frequently employed in modelling of bacterial growth. We will use this form in Section Part 3 to estimate the growth parameters from experimental data for each type grown in pure culture.

### Simulation method

Continuous-time birth processes can in principle be simulated using the Gillespie algorithm (Gillespie, 1976, 1977). This is an exact procedure for the production of sample paths. However, carrying out the Gillespie algorithm becomes time consuming when the populations are large. This is because the number of events occurring in the population per unit time is proportional to the size of the population. Bacterial populations often contain numbers of individuals of order $10^9$; in the case of our simulations population sizes can go up to $5.71 \times 10^8$. This makes continuous-time Gillespie simulations unrealistic.

In such situations one therefore has to resort to approximations. One such approximation method is the so-called $\tau$-leaping variant of Gillespie's algorithm (Gillespie, 2001). This approach proceeds in discrete time and assumes that total event rates in the population are constant over each step. The number of offspring produced in each step is then a Poissonian random number. Even in the case of constant reproduction rates $\lambda_i$ this is no longer an exact procedure. Mathematically, the sampling is not from the true solution of the master equation [the negative binomial distribution in Eq. (6)]. This is because the $\tau$-leaping algorithm does not capture events in which an offspring generated in one step undergoes a further reproduction event in the time step, a sequence of processes which is in principle possible in the continuous-time model.

Like the $\tau$-leaping method, our simulations proceed in discrete time, with a time step $\Delta t$. However we do not make the above Poissonian approximation, instead our sampling is from the negative binomial distribution [Eq. (6)]. For a pure birth process this would mean to sample the number of individuals $n_i = n_i(t + \Delta t)$ at time $t + \Delta t$ from the distribution

$$p_{n_i}(t + \Delta t; n_i(t)) = \binom{n_i - 1}{n_i - n_i(t)} e^{-\lambda_i n_i(t)\Delta t} \left(1 - e^{-\lambda_i \Delta t}\right)^{n_i - n_i(t)}, \tag{10}$$

where $n_i(t)$ is the number of individuals of strain $i$ at time $t$, and where $\lambda_i$ is the reproduction rate for this step. As discussed above and unlike $\tau$-leaping, this is an exact procedure if the rate $\lambda_i$ does not vary over the time step.

In our model the reproduction rates $\lambda_i$ depend on the total number of individuals in the population, $n_T$. They therefore become time dependent, with interactions between the strains. The simulation method is then an approximation of the continuous-time process. We provide a comparison against simulations of the full continuous-time model further below.

#### Mutations

The simulation model described in the conceptual diagram in Figure 3 of the main text involves mutations. These can occur from sensitive type bacteria $S$ to either types $R$ or $N$, and, in turn, from type $R$ and type $N$ to type $D$. The model excludes the possibility of direct mutations from $S$ (sensitive) to $D$ (double resistance).

In order to incorporate mutations, we first calculate the number of offspring produced by a strain in a time step. Starting with $n_i^0 = n_i(t)$ cells of type $i$ at the beginning of the time step we write $n_i(t + \Delta t)$ for the number of cells of strain $i$ at the end of the step. The number of offspring $m_i$ generated in this step is then

$$m_i = n_i(t + \Delta t) - n_i^0. \tag{11}$$

Therefore, the probability $P_{m_i}(t + \Delta t; n_i^0)$ that $m_i$ offspring have been produced in the time step from $t$ to $t + \Delta t$ is equal to the probability that there are $n_i(t) = n_i^0 + m_i$ individuals of strain $i$ in the population at time $t + \Delta t$.

Using Eq. (6) this results in the following expression for the distribution of $m_i$:

$$P_{m_i} = \binom{n_i^0 + m_i - 1}{m_i} p_i^{n_i^0} q_i^{m_i}. \tag{12}$$

We assume that offspring of an individual of type $S$ are of type $R$ with probability $\mu_R$ or of type $N$ with probability $\mu_N$. With the remaining probability $1 - \mu_R - \mu_N$ the offspring of a parent of type $S$ is also of type $S$ (no mutation). Similarly, an offspring of a parent of type $R$ is of type $D$ with probability $\mu_N$, and an offspring of a parent of type $N$ is of type $D$ with probability $\mu_R$.

Summarising, the simulations proceed as follows:

1. Assuming that there are $n_i(t)$ individuals of strains $i \in \{S, R, N, D\}$ at time $t$, compute $\lambda_i$ via Eq. (7), and from this the $p_i = e^{-\lambda_i \Delta t}$ and $q_i = 1 - p_i$.

2. Then draw the number of offspring $m_i$ for each type from the distribution in Eq. (12).

3. We first focus on the number offspring $m_S(t)$. The number of these that experience a mutation into type $R$ is binomially distributed $n_{SR}(t) =$ Binomial$[m_S(t), \mu_R]$. The number of offspring with mutations into type $N$ follow $n_{SN}(t) =$ Binomial$[m_S(t), \mu_N]$. The remainer of the offspring of a parent of type $S$ is of type $S$, $n_{SS}(t) = m_S(t) - n_{SR}(t) - n_{SN}(t)$. Similarly, the number of offspring of $R$ with a mutation into type $D$ will be $n_{RD}(t) =$ Binomial$[m_R(t), \mu_N]$ and the remainder $n_{RR}(t) = m_R(t) - n_{RD}(t)$ is type $R$. Analogously, for strain $N$ we have $n_{ND}(t) =$ Binomial$[m_N(t), \mu_R]$ and $n_{NN}(t) = m_N(t) - n_{ND}$ for offspring of type $D$ and $N$, respectively.

   The number of individuals at the end of the time-step are then

$$\begin{aligned} n_S(t + \Delta t) &= n_S(t) + n_{SS}(t), \\ n_R(t + \Delta t) &= n_R(t) + n_{RR}(t) + n_{SR}(t), \\ n_N(t + \Delta t) &= n_N(t) + n_{NN}(t) + n_{SN}(t), \\ n_D(t + \Delta t) &= n_D(t) + m_D(t) + n_{RD}(t) + n_{ND}(t). \end{aligned} \tag{13}$$

4. Increment time by $\Delta t$ and go to step 1.

We have written Binomial$[n, p]$ for a binomial random number with parameters $n$ and $p$; the probability that such a random number takes value $j$ is $p_j = \frac{n!}{j!(n-j)!} p^j (1-p)^{n-j}$, for $j = 0, 1, \ldots, n$.

The periodic dilution in simulations of the laboratory experiments is implemented separately, for futher details see Section Part 3.

#### Estimating growth parameters from experimental data

We used experimental data to estimate values for the parameters $r_i$ and $k_i$ introduced in Eq. (7). We carried out growth curve experiments of the wild-type sensitive strain (type $S$) and resistant strains (types $R$, $N$ or $D$) by measuring optical density (OD) over time (Supplementary

Figure 3A), under conditions matching the selection experiment (see Methods in the main text). For each resistant type, we isolated five independent mutant clones derived from the wild-type $S$ type via independent fluctuation tests (see Methods in the main text).

The growth curves deviate from a typical 'S'-shaped logistic curve, instead characterised by two 'plateaus', reminiscent of diauxic growth. Diauxic growth has been previously reported for MH media (CAVALLERO *et al.*, 1990), as well as for other rich media (WANG and KOCH, 1978; SEZONOV *et al.*, 2007). We account for this two-stage growth by fitting the data to growth curves of the form

$$n_i(t) = n_i^{(1)}(t)H(t_c - t) + n_i^{(2)}(t)H(t - t_c), \tag{14}$$

where $n_i^{(1)}(t)$ and $n_i^{(2)}(t)$ are the same functionals as Eq. (9):

$$n_i^{(1)}(t) = \frac{k_i^{(1)} n_i^c}{n_i^c + [k_i^{(1)} - n_i^c]e^{-r_i^{(1)}(t-t_i^c)}}, \quad n_i^{(2)}(t) = \frac{k_i^{(2)} n_i^c}{n_i^c + [k_i^{(2)} - n_i^c]e^{-r_i^{(2)}(t-t_i^c)}}, \tag{15}$$

and where $H(t)$ is the Heaviside step function, $H(t) = 0$ for $t < 0$, $H(t = 0) = 1/2$ and $H(t) = 1$ for $t > 0$. $k_i^{(1)}$ and $k_i^{(2)}$ are the carrying capacities of strain $i$ before and after the switch. The functions $n_i^{(1)}(t)$ and $n_i^{(2)}(t)$ are logistic growth curves that represent the two regimes of the diauxic growth. They are solutions of the one-strain logistic equation, Eq. (9). The growth switches from $n_i^{(1)}$ to $n_i^{(2)}$ at time $t_c$, where $n_i^{(1)}(t)$ describes the growth for $t < t_i^c$ and $n_i^{(2)}(t)$ for $t > t_i^c$. Using the definition of the Heaviside function $H(t = 0) = 1/2$, one has $n_i(t) = n_i^c$ at $t = t_i^c$.

We illustrate the curve fitting for the rifampicin resistant type grown in the presence of the rifampicin, at the same concentrations used in the selection experiment (Supplementary Figure 4). For the purposes of the fit we only considered the first 25 h of the growth experiment, to eliminate potential complications introduced by evaporation of the growth medium. We measure time in units of hours, and the growth rates $r_i$ are thus expressed in units of $h^{-1}$. The fits were performed using MATLAB 2016a, with its Non-linear Least Squares fitting method (code provided, see Data Availability statement). Typically, these parameters explain a very large majority of the variation in OD (Supplementary Table 6), except where there is effectively no growth (e.g. of $S$, $R$ and $N$ strains in high concentrations of the antibiotic combination).

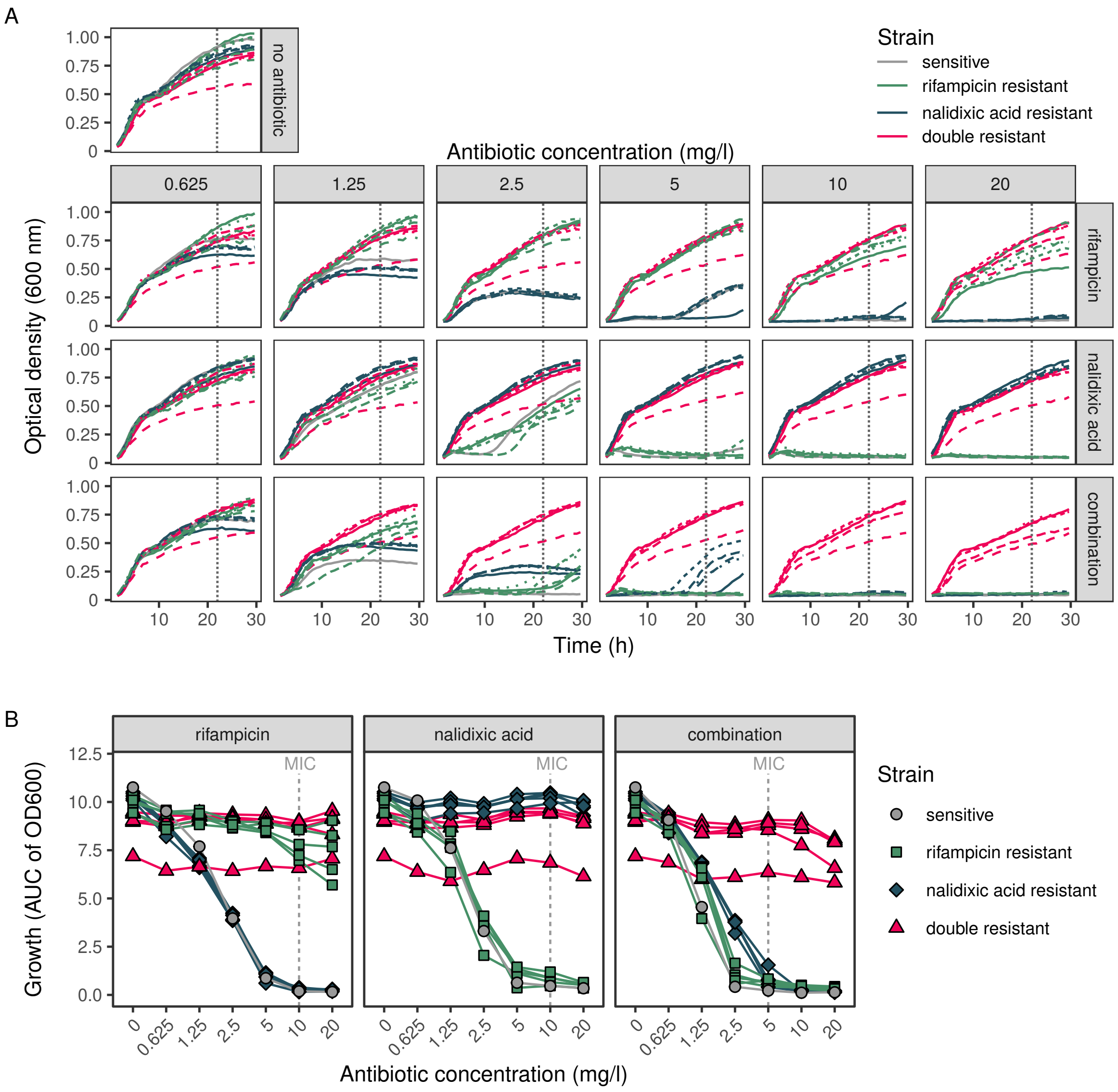


**Supplementary Figure 3:** Growth of sensitive, single-, and double-drug resistant strains under antibiotic concentrations used during the selection regime. Strains shown here were derived from the wild-type background, independently of the selection experiment. A. Growth curves, with each line depicting one of five different strains (indicated by line type), averaged over replicate growth curves ($n = 5$). Note curve shapes reminiscent of diauxic growth. B. Area under the curve for each strain.

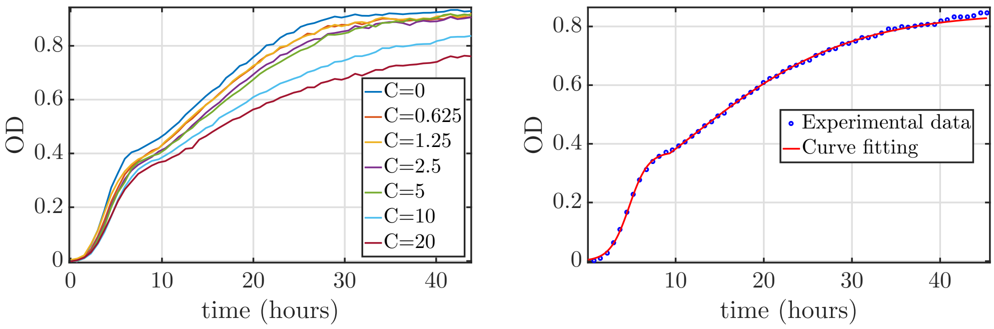


**(a)** **(b)**

**Supplementary Figure 4:** A. Example growth curves of a rifampicin-resistant clone under rifampicin treatment, for several drug concentrations, $C$ (mg/l). B. Curve fitting for $C = 10$ mg/l of the rifampicin treatment for the rifampicin-resistant strain, resulting in estimates $k_R^{(1)} = 0.372689$ [95% CI: (0.3652, 0.3801)], $r_R^{(1)} = 0.921300$ [95% CI: (0.8678, 0.9748)], $k_R^{(2)} = 0.848618$ [95% CI: (0.8038, 0.8935)], $r_R^{(2)} = 0.110723$ [95% CI: (0.0998, 0.1216)], $R_R^2 = 0.99966$. Parameter estimates and $R^2$ for the other curves are shown in Supplementary Figure 6 and Supplementary Table 6, respectively. OD–optical density.

**Supplementary Table 6:** $R^2$ values obtained from fitting Eq. (15) to the data in Supplementary Figure 3A using the approach illustrated in Supplementary Figure 4 for the strains $S$, $R$, $N$, and $D$.

| Treatment | Transfer | Concentration (mg/l) | $R_R^2$ | $R_N^2$ | $R_D^2$ | $R_S^2$ |
|---|---|---|---|---|---|---|
| no antibiotic | 1–6 | 0.31250 | 0.99957 | 0.99925 | 0.99907 | 0.99957 |
| rifampicin | 1 | 0.625 | 0.99960 | 0.99738 | 0.99966 | 0.99956 |
| rifampicin | 2 | 1.25 | 0.99964 | 0.99625 | 0.99958 | 0.99244 |
| rifampicin | 3 | 2.50 | 0.99958 | 0.98607 | 0.99946 | 0.99552 |
| rifampicin | 4 | 5 | 0.99956 | 0.99822 | 0.99967 | 0.99947 |
| rifampicin | 5 | 10 | 0.99966 | 0.99725 | 0.99969 | 0.94550 |
| rifampicin | 6 | 20 | 0.99933 | 0.99165 | 0.99974 | 0.37149 |
| nalidixic acid | 1 | 0.625 | 0.99959 | 0.99955 | 0.99953 | 0.99945 |
| nalidixic acid | 2 | 1.25 | 0.99974 | 0.99953 | 0.99964 | 0.99948 |
| nalidixic acid | 3 | 2.50 | 0.99763 | 0.99957 | 0.99944 | 0.99309 |
| nalidixic acid | 4 | 5 | 0.98397 | 0.99958 | 0.99970 | 0.98801 |
| nalidixic acid | 5 | 10 | 0.97461 | 0.99949 | 0.99970 | 0.96445 |
| nalidixic acid | 6 | 20 | 0.96781 | 0.99970 | 0.99932 | 0.95763 |
| combination | 1 | 0.625 | 0.99948 | 0.99940 | 0.99950 | 0.99913 |
| combination | 2 | 1.25 | 0.99962 | 0.99939 | 0.99964 | 0.99924 |
| combination | 3 | 2.50 | 0.97307 | 0.99931 | 0.99963 | 0.95726 |
| combination | 4 | 5 | 0.97735 | 0.98672 | 0.99960 | 0.88433 |
| combination | 5 | 10 | 0.93615 | 0.98740 | 0.99955 | 0.19791 |
| combination | 6 | 20 | 0.97295 | 0.97101 | 0.99955 | 0.35897 |

In order to include the diauxic-like behaviour in the simulation model, we switch between the two growth regimes when the total population size $n_T(t)$ reaches a threshold $n_T^c$. This threshold is related to the $n_i^c$ obtained from the fits of single-strain growth experiments to the diauxic-like behaviour in Eq. (14) as we will explain next.

The time $t_i^c$ and population size $n_i^c$ at which the growth switches from $n_i^{(1)}(t)$ to $n_i^{(2)}(t)$ is different for every strain, treatment, and drug concentration considered. Note that these parameters are obtained from growth experiments with strains grown on their own, whereas multiple strains are present in the simulation model. We therefore use the $n_i^c$ distributions to choose a single time point at which to switch regimes for the entire population based on the procedure outlined below. Supplementary Figure 5 shows the distribution of $n^c$, the switching population values, obtained for the different treatments. Each graph shows a histogram of the values, $n^c$, obtained from the different drug concentrations and strains for the given treatment. Since there are 6 drug concentrations as each growth experiment proceeds, and $4$ different strains, this results in $24$ values for $n^c$ per treatment. Evidently, for experiments in which no drug is given we only obtain four values of $n^c$ (one for each strain). We do not show the corresponding histogram.

Each of the histograms is bimodal, with one peak close to $n^c = 0$, and the other at $n^c \approx 0.3$ on the optical-density scale. The left peak at $n^c \approx 0$ corresponds to strains incapable

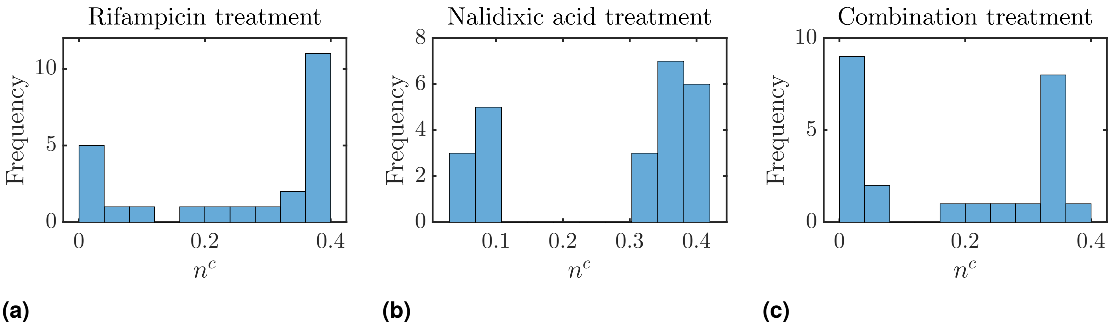


**Supplementary Figure 5:** Histograms of $n^c$ obtained from fitting the two-stage 'diauxic-like' growth curves (Eqs. 14 and 15) to experimental data; rifampicin, nalidixic acid, combination treatments in panels A, B and C, respectively. Each graph includes all the values (24 in total) of $n_i^c$ obtained for each drug concentration and strain.

**Supplementary Table 7:** Switching value, $n_T^c$, for the different treatments.

| Treatment | Switching $n_T^c$ |
|---|---|
| No antibiotic | 0.4108 |
| Rifampicin | 0.3417 |
| Nalidixic acid | 0.3709 |
| Combination | 0.3154 |

of growing in those environments (e.g. strain $S$ for high values of antibiotic concentration, $C$). Taking the average across this distribution would therefore not be representative of growing strains. Instead, we estimate $n_T^c$ as the mean value obtained from the peak on the right in each histogram, i.e. cases where strains were capable of growing. Results are shown in Supplementary Table 7. The value obtained in this way differs between the different treatments. For the case of no antibiotic treatment we use the average over the four strains as threshold value $n_T^c$.

Parameters estimated in this way are given in Supplementary Figure 6. As these parameters have been estimated from OD growth curves, they have so far been expressed in units of OD. In the next section, we will describe the relationship between OD and the number of bacteria.

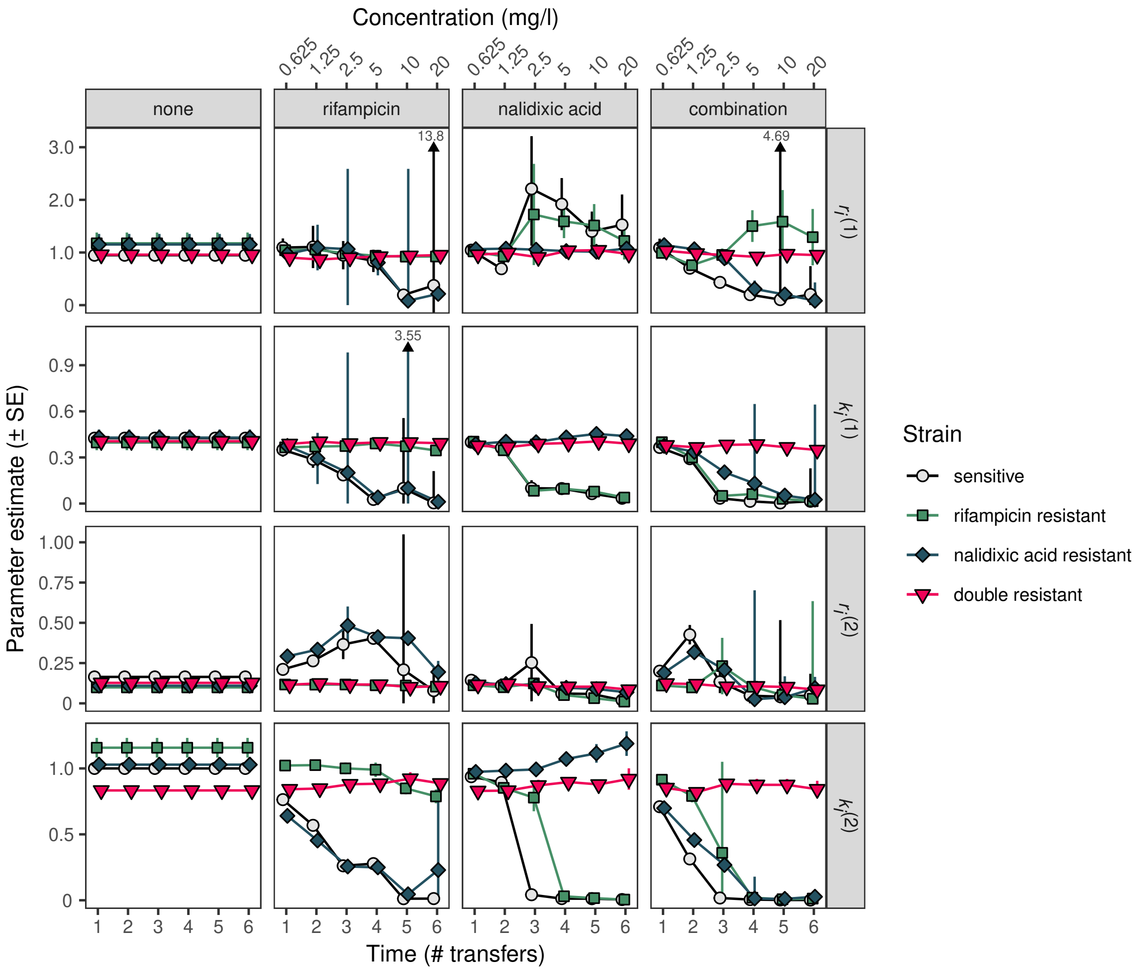


**Supplementary Figure 6:** Parameter estimates for $k_i^{(j)}$ and $r_i^{(j)}$ from growth curve fitting used in simulations (See Supplementary Figure 3A and Supplementary Figure 4). Error bars represent the standard error of the estimate across five independent strains (upper bounds of extreme error bars have been truncated as indicated by an arrow and its value; lower bounds are truncated at zero). Points have been displaced on the horizontal axis for clarity.

#### Relationship between optical density and bacterial population size

To simulate population dynamics, the parameters we have estimated need to be expressed in terms of numbers of bacteria, rather than in units of OD. We therefore determined the relationship between OD and the number of bacteria present. Following growth in liquid MH (with and without antibiotics, as in the selection experiment), serial dilutions were plated on 90 mm Petri dishes containing LB agar (without antibiotics), and colony forming units (CFUs) were counted following overnight growth. We compared the fit of a linear ($y = mx + b$) and non-linear ($y = mx^a + b$) relationship between $\log(\text{CFUs})$ and $\log(\text{OD})$. Both models were fit using Bayesian non-linear regression using `brm()` from the `brms` package (Bürkner, 2017, 2018) in R (R Core Team, 2019). Model comparison was performed using the expected log pointwise predictive density calculated (ELPD) using leave-one-out (PSIS-LOO) cross-validation with `loo_compare()` (Vehtari *et al.*, 2017). The linear and non-linear models had equivalent predictive power when corrected for the number of parameters fit, therefore the simpler, linear, model was used. Parameter estimates for both models are shown below in Supplementary Table 8. The linear model and the underlying data are plotted in Supplementary Figure 7A.

We also performed kill curves to quantify killing that occurs when populations first experience antibiotic treatment (Supplementary Figure 7B).This was performed by pin replicating an overnight culture (1/200 dilution) of the sensitive ancestor into antibiotic containing medium (0.625, 1.25, 2.5, 5, 10, 20 mg/l of each antibiotic treatment). Population sizes were estimated by plating dilutions on LB agar after 2, 4, 6, 8, and 24 h of exposure.

**Supplementary Table 8:** Bayesian non-linear regression parameters describing two possible relationships between optical density (OD) and colony forming units (CFU), as estimated using `brm()` from the brms package. Both models had equivalent predictive power as measured by ELPD using the LOO cross-validation method.

Regression model of the form CFU = $m$OD + $b_{\text{overall}} + b_i$

| | Estimate | Standard error |
|---|---|---|
| $\text{ELPD}_{\text{LOO}}$ | -125.0 | 14.5 |
| $\text{p}_{\text{LOO}}$ | 8.4 | 1.2 |
| LOOIC | 250.0 | 29.1 |

| Parameter | Estimate | Estimate error | 95% CI |
|---|---|---|---|
| $m$ | 1.895 | 0.065 | $(1.772, 2.026)$ |
| $b_{\text{none}}$ | 3.297 | 0.147 | $(3.005, 3.581)$ |
| $b_{\text{rifampicin}}$ | -0.302 | 0.099 | $(-0.499, -0.109)$ |
| $b_{\text{nalidixic acid}}$ | -0.345 | 0.112 | $(-0.565, -0.123)$ |
| $b_{\text{combination}}$ | -0.197 | 0.109 | $(-0.415, 0.018)$ |
| $b_{\text{none:concentration}}$ | -0.010 | 0.988 | $(-1.956, 1.926)$ |
| $b_{\text{rifampicin:concentration}}$ | 0.286 | 0.080 | $(0.132, 0.443)$ |
| $b_{\text{nalidixic acid:concentration}}$ | 0.187 | 0.087 | $(0.022, 0.364)$ |
| $b_{\text{combination:concentration}}$ | 0.098 | 0.088 | $(-0.072, 0.268)$ |

Regression model of the form CFU = $m\text{OD}^a$ + $b_{\text{overall}} + b_i$

| | Estimate | Standard error |
|---|---|---|
| $\text{ELPD}_{\text{LOO}}$ | -123.9 | 15.2 |
| $\text{p}_{\text{LOO}}$ | 9.2 | 1.5 |
| LOOIC | 247.7 | 30.3 |

| Parameter | Estimate | Estimate error | 95% CI |
|---|---|---|---|
| $m$ | 2.990 | 0.537 | $(1.990, 4.069)$ |
| $a$ | 0.762 | 0.093 | $(0.607, 0.968)$ |
| $b_{\text{none}}$ | 2.081 | 0.600 | $(0.906, 3.231)$ |
| $b_{\text{rifampicin}}$ | -0.281 | 0.100 | $(-0.475, -0.086)$ |
| $b_{\text{nalidixic acid}}$ | -0.316 | 0.112 | $(-0.532, -0.092)$ |
| $b_{\text{combination}}$ | -0.168 | 0.107 | $(-0.385, 0.038)$ |
| $b_{\text{none:concentration}}$ | 0.012 | 0.991 | $(-1.918, 1.970)$ |
| $b_{\text{rifampicin:concentration}}$ | 0.282 | 0.079 | $(0.126, 0.432)$ |
| $b_{\text{nalidixic acid:concentration}}$ | 0.169 | 0.087 | $(0.001, 0.341)$ |
| $b_{\text{combination:concentration}}$ | 0.084 | 0.084 | $(-0.080, 0.253)$ |

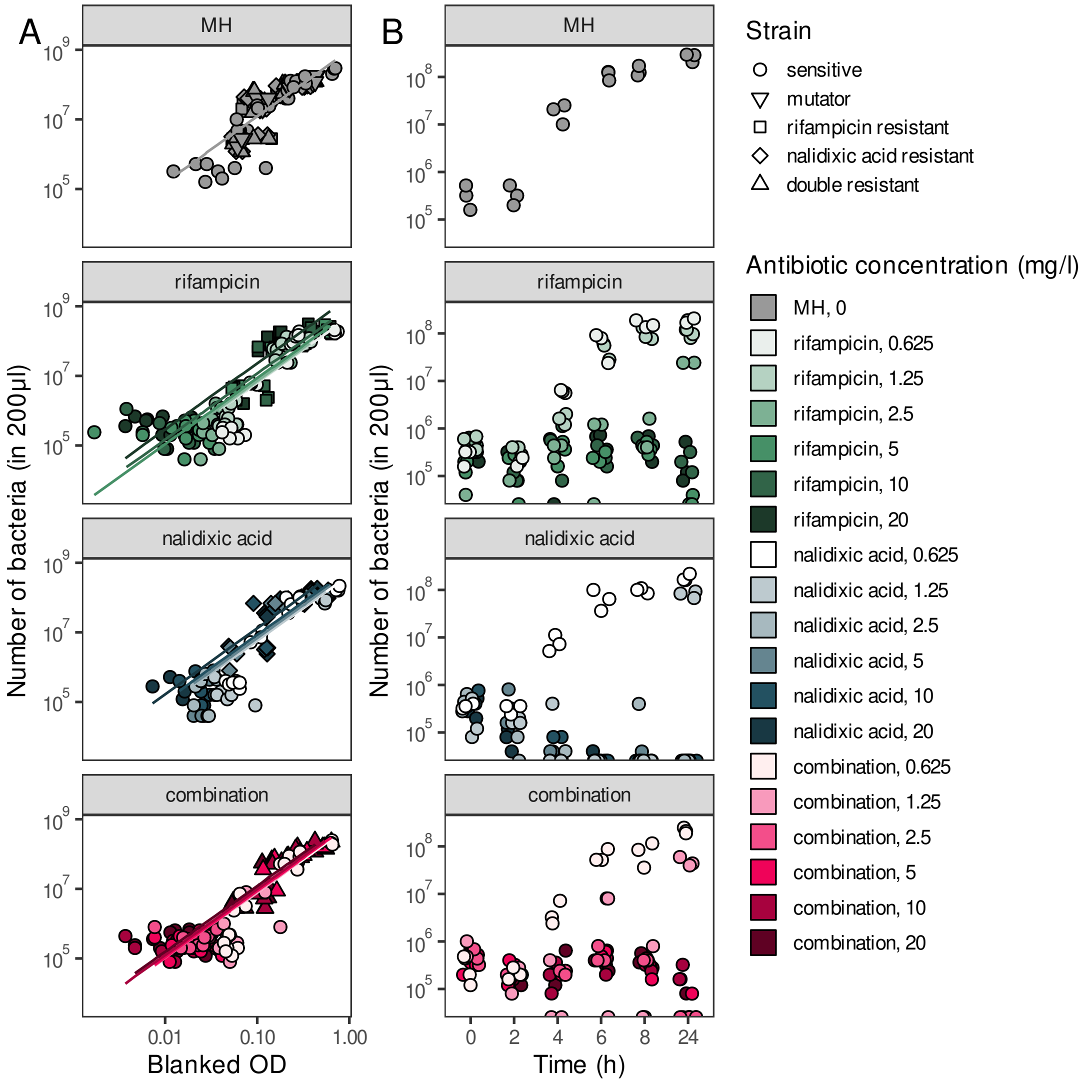


**Supplementary Figure 7:** A. Relationship between blank corrected optical density (Blanked OD) and number of bacteria (measured by colony forming units) in antibiotic concentrations experienced during the selection experiment. Regression lines are the best fitting linear model shown in Supplementary Table 8. B. Time series of sensitive bacterial populations exposed to antibiotic concentrations experienced during the selection experiment.

### Simulating the experiments

Simulation conditions are given in Supplementary Table 9. We use the values $k_i^{(1)}, r_i^{(1)}, k_i^{(2)}, r_i^{(2)}$ of each strain and treatment, and $n_T^c$ obtained as defined above, for simulations of the stochastic model. Simulations are started from an initial population of $5.71 \times 10^6$ sensitive bacteria (type $S$); a frequency $u$ of these are mutators and the remaining fraction $(1-u)$ consists of wild-type bacteria. In the simulations we use $\mu_R = 6.7 \times 10^{-9}$ and $\mu_N = 7.4 \times 10^{-10}$, as motivated in the main text. Mutators have a 80-fold increased mutation rates, $\mu_R$ and $\mu_N$, compared to the wildtype.

**Supplementary Table 9:** Simulation conditions for resistance evolution simulations.

| Parameter | Value |
|---|---|
| Replicates | 1000 |
| Initial population size | $5.71 \times 10^6$ |
| Mean proportion transferred during dilution step | 0.005 |
| Duration between dilution steps | 22 h |
| Length of time per time-step ($\Delta t$) | 0.01 h |
| Mutation rate to rifampicin resistance ($\mu_R$) | $6.7 \times 10^{-9}$ mutations/replication |
| Mutation rate to nalidixic acid resistance ($\mu_N$) | $7.4 \times 10^{-10}$ mutations/replication |
| Mutator effect | 80-fold increase |

The simulated experiment consists of six days of growth, where the concentration of antibiotic(s) is initially 0.625 on day 1, and doubled on each of the days 2 to 6. This procedure was chosen to reflect the experimental conditions. The length of each time-step of the simulation, $\Delta t$, is expressed in units of hours (h). It has to be sufficiently small so that the simulations maintain precision and capture the behaviour of the original continuous-time model. We have set $\Delta t = 0.01$ h, which is much smaller than the characteristic time of *E. coli* division, which is approximately 0.3–0.6 h (Helmstetter and Cooper, 1968). this choice approximates simulations of the full model in continuous time to a good accuracy, while leading to a considerable reduction of the simulation time. In Supplementary Figure 8, we compare the distributions of the number of individuals of each strain obtained from these discrete-time simulations with simulations of the original continuous-time model. The discrete-time simulations approximate the original model with good accuracy. For the simulation conditions used in this figure, the simulation time of one single run (i.e. 22 h of the experiment) is approximately 500 times faster with our approach relative to the Gillespie algorithm on the same computing hardware.

An important aspect to take into account is the dilution carried out in the laboratory experiments at the end of each day, when $1/200$ of the population is carried forward to the next day, and the rest discarded. In the evolution literature, this is referred to as a population bottleneck. In simulations we cycle through all members of the population at the end of each (simulated) day and retain each individual with probability $1/200$; otherwise the individual is

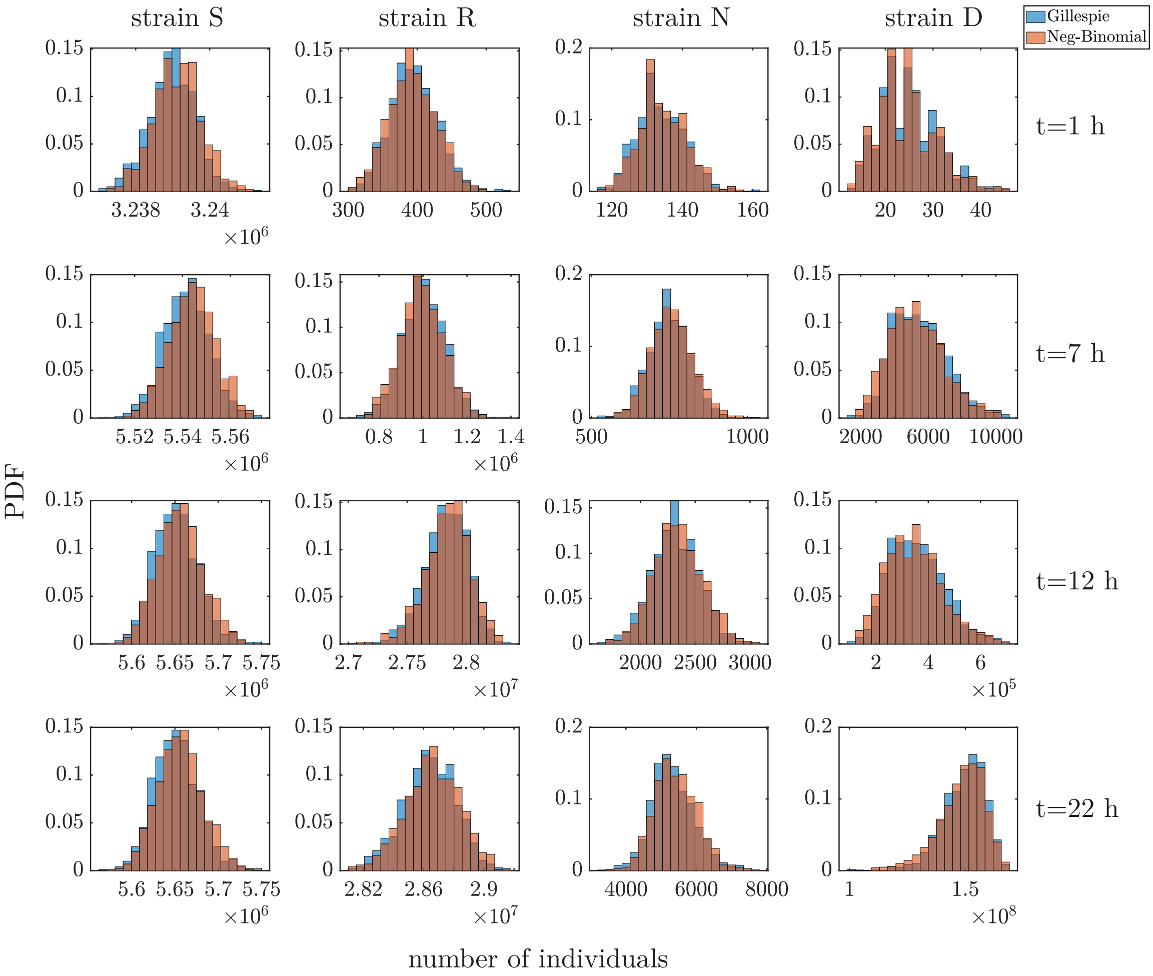


**Supplementary Figure 8:** Distributions of the number of individuals of each strain at different times obtained from continuous-time simulations (Gillespie algorithm) and discrete-time simulations (based on the negative binomial distribution, see Section Part 3). Data from simulations of the continuous-time model are shown as blue bars, those from the discrete-time approach as light orange bars. Dark brown colour indicates an overlap of both types of bars. The simulation parameters used are the parameter estimates from day 4 of the combination treatment (see Supplementary Figure 6). We set an initial number of wild-type individuals equal to $n_S = 2.8 \times 10^6$, $n_R = 100$, $n_N = 100$, and $n_D = 10$. For the discrete-time approach we set $\Delta t = 0.01$. The distributions were constructed over 1000 runs.

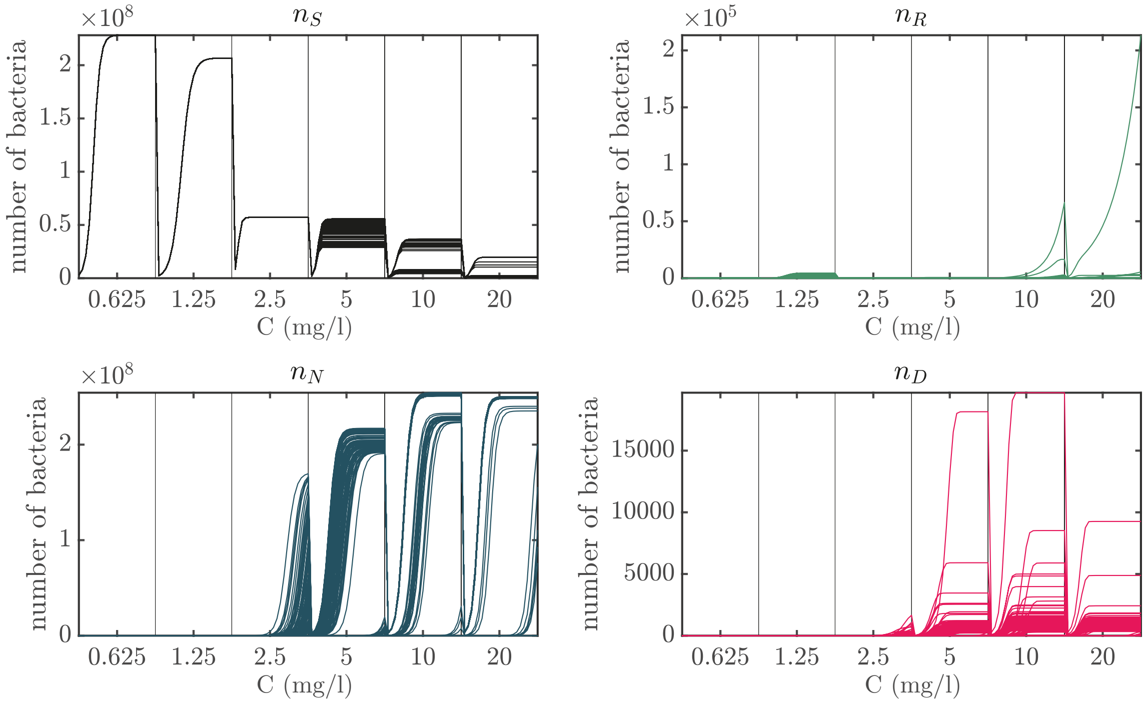


**Supplementary Figure 9:** Simulated growth curves for nalidixic acid treatment. Each panel shows 100 independent simulation runs. The data shown aggregates wildtype and mutator bacteria. $C$–antibiotic concentration (mg/l). Initial condition: $5.71 \times 10^6$ sensitive bacteria, with a frequency $u = 0.3$ of mutators. Curves for each value of $C$ were simulated over 22 h. We have used $\Delta t = 0.01$ h. Compare to Extended Data Figure 1D (nalidixic acid column).

removed. This represents an independent Bernoulli event on the level of individual bacteria. The number of each type $i$ carried forward follows a binomial distribution with mean $\mu = n_i/200$ and variance $\sigma^2 = n_i \times 1/200 \times 199/200 = n_i \times 4.975 \times 10^{-3}$, where $n_i$ is the number of type $i$ individuals. This allows the possibility that a strain type goes extinct if it arises too late into the growth cycle to achieve appreciable frequency. An example of the growth curves obtained from the simulations, including the dilution step, is presented in Supplementary Figure 9.

Supplementary Figure 14 shows the dynamics of mutations spreading within four example populations exposed to the combination treatment (equivalent to the main text Figure 4A). The interquartile range across all replicate simulations is presented in Supplementary Figure 15 (equivalent to the main text Figure 4).

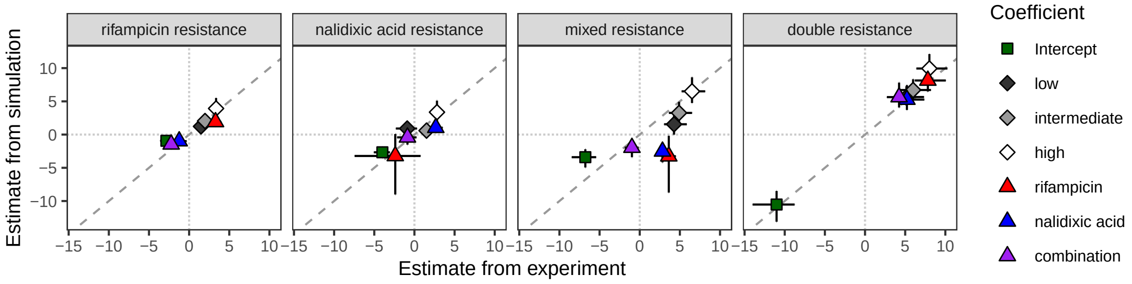


**Supplementary Figure 10:** Comparison of coefficients from Bayesian categorical models fitted to simulation and experimental data (Models M1 and M4, respectively). Dashed diagonal line indicates 1:1 line; points falling on this line indicate coefficients are equal in both simulation and experiment.

##### Comparison between simulation and experimental results

To determine concordance between simulations and experimental results, we fit a Bayesian categorical model to the simulation results and compared the estimated coefficients. The same statistical model and priors were used to analyse the simulation data as were used in the analysis of the experimental results, with the exception of an absence of random effects of 'position' (see section Part 1). As for the experimental results, the full model with the interaction did not provide a better fit than the main effects model (WAIC $1486.5 \pm 52.5$ SE vs. $1515.0 \pm 53.1$ SE). The estimated coefficient obtained from fitting the main-effects model to the output from the simulations are therefore given in Supplementary Table 10. Estimated coefficients from the simulation closely matched those from the experiment (Supplementary Figure 10), with the exception of the 'mixed resistance' category, which the tended to estimate a different sign for the effects of the nalidixic acid and combination treatments.

#### Other simulation conditions

##### Simulating different antibiotic dose escalation regimes

In our experiments and main set of simulations, we have considered a particular dose escalation regime. Here we use the simulation model to predict whether other dose escalation regimes would affect the role of mutators in multi-resistance evolution. For example, if the concentration of the combination treatment increases faster, the sensitive and single-resistant strains will have less opportunity to acquire sequential resistance mutations (or conversely, if the concentration rises more slowly, there will be more opportunity for multi-resistance). To simulate reaching MIC at an earlier time, we time-shifted the parameter values relative to the original simulations (e.g. for MIC occurring on the third transfer instead of the fifth, the new values for transfers 1–4 corresponded to the parameters estimated for the 3rd, 4th, 5th (MIC), and 6th

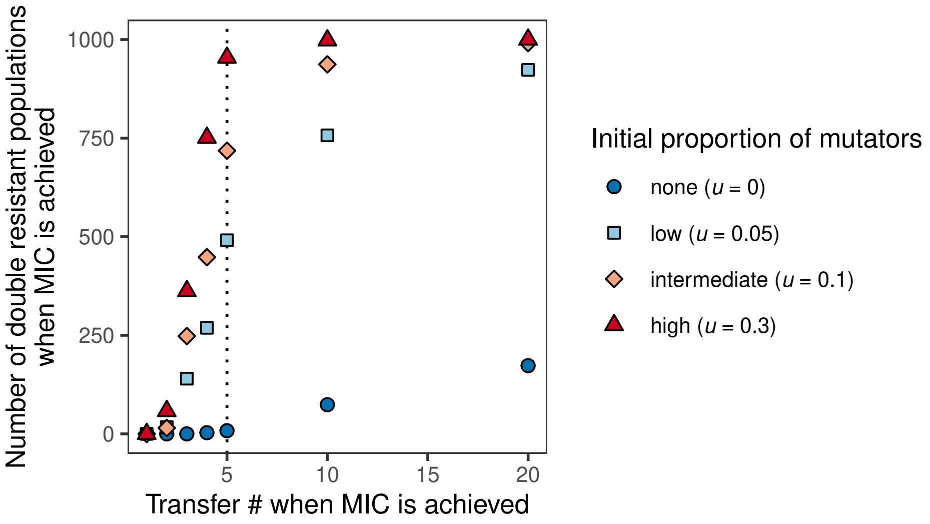


**Supplementary Figure 11:** Simulations of combination treatment where MIC is achieved after different numbers of population transfers ($n = 1000$ replicate simulations). Dotted vertical line corresponds to the simulation conditions used in the main text Figures 3 and 4

transfers of the original simulations). For simulations with MIC occurring later than the original simulations, the parameter values were 'stretched' (e.g. for MIC occurring on the tenth transfer instead of the fifth, the values used for transfers 1–10 corresponded to the original parameters estimated for the 1st, 1st, 2nd, 2nd, 3rd, 3rd, 4th, 4th, 5th, 5th, 6th and 6th transfers).

The overall proportion of multi-resistant populations increases with the time taken to reach MIC. When MIC was achieved earlier, there is overall less multi-resistance, but it nevertheless occurs for the highest mutator frequency, even when achieved just before the first transfer. When MIC was achieved later, the model predicts that a proportion of purely wild-type populations should achieve multi-resistance when the duration of the experiment is doubled or quadrupled (6.4% or 18.9% of populations, respectively), but that a small proportion of mutators still results in vastly higher multi-resistance (76.3% and 92.2% of populations, respectively). While multi-resistance is reduced when MIC occurs sooner, populations with mutators can still evolve multi-resistance, even for very fast dose escalation.

#### Simulating strains with a cost of resistance and logistic growth

Our simulations used parameter values estimated from growth curve data from resistant strains, and a two-stage population growth model to match the shape of growth curves under our environmental conditions (rather than logistic growth, as is typically used to model microbial growth). The estimated parameters did not always demonstrate a 'cost of resistance' (Supplementary Figure 6), i.e. the phenomenon that resistance to an antibiotic tends to be associated with a fitness deficit in environments where that antibiotic is not present (Andersson, 2006, though resistance need not be costly, Lenormand *et al.* 2018). This was particularly evident in the antibiotic-free environment where single-resistant strains had a higher growth rate ($r_i^{(1)}$) than the sensitive strain (Supplementary Figure 6). We therefore tested whether we would observe similar patterns of resistance emergence if we imposed arbitrary growth parameters for single resistant types $A$ and $B$ and double resistant type $D$ that reflect a cost of resistance and standard logistic growth (keeping other model parameters, e.g. mutation rate, the same). Growth parameters were chosen so that only strains that would be expected to have a benefit in a given environment actually do (i.e. type $A$ has an advantage in environments with drug A, type $B$ with drug $B$, Supplementary Figure 12A). We reduced our two-stage model to a logistic growth model by setting $r_i^{(1)} = r_i^{(2)}$ and $k_i^{(1)} = k_i^{(2)}$.

Using these parameters, we again observe that mutators allow multi-resistance to arise in both single-drug and combination treatment environments (Supplementary Figure 12B), as we observed in the original simulations (Figure 3). Hence, the patterns of resistance emergence we observed are not specific to the parameter estimates obtained from our experiments. Moreover, within populations, the spread of double resistance follows the hitch-hiking of a mutator allele along with a single resistance mutation, which subsequently increases the mutation rate of the population sufficiently to observe the secondary resistance mutation (Supplementary Figure 12C–F). This matches the same trend as the original simulations (Figure 4 and Supplementary Figure 15).

#### Simulating very large population sizes

Our simulations have shown that mutators contribute to multi-resistance evolution because they increase the supply of resistance mutations, allowing independent resistance mutations to be acquired sequentially. However, mutational supply also depends on population size, which raises the possibility that multi-resistance may also evolve in large populations without mutators. We used our simulation model to determine how large a purely wild-type population would need to be to observe multi-resistance without mutators. Supplementary Figure 13 shows the average proportion of each type of strain under different maximum population sizes after six days of simulated evolution. Multi-resistance reached appreciable frequencies for maximum population sizes between $5.71 \times 10^{10}$ (combination treatment) and $5.71 \times 10^{13}$ (no antibiotic treatment), which in all cases is orders of magnitude larger than the population size of $5.71 \times 10^{8}$ in our experimental conditions. Bacterial population sizes in infections are typically smaller than this range (Orenstein and Wong, 1999; Opota *et al.*, 2015), although not always (Bingen *et al.*, 1990; Stressmann *et al.*, 2011). Commensal populations, however, can achieve population sizes in the range of $10^{10}$ or more (Savage, 1977; Mason and Richardson, 1981), and $10^{13}$ bacteria is comparable in size to the entire human-associated microbiome (Sender *et al.*, 2016). Thus, multi-resistance may also be found without mutators in the largest microbial populations (although these populations also often contain mutators, Ramiro *et al.*, 2020).

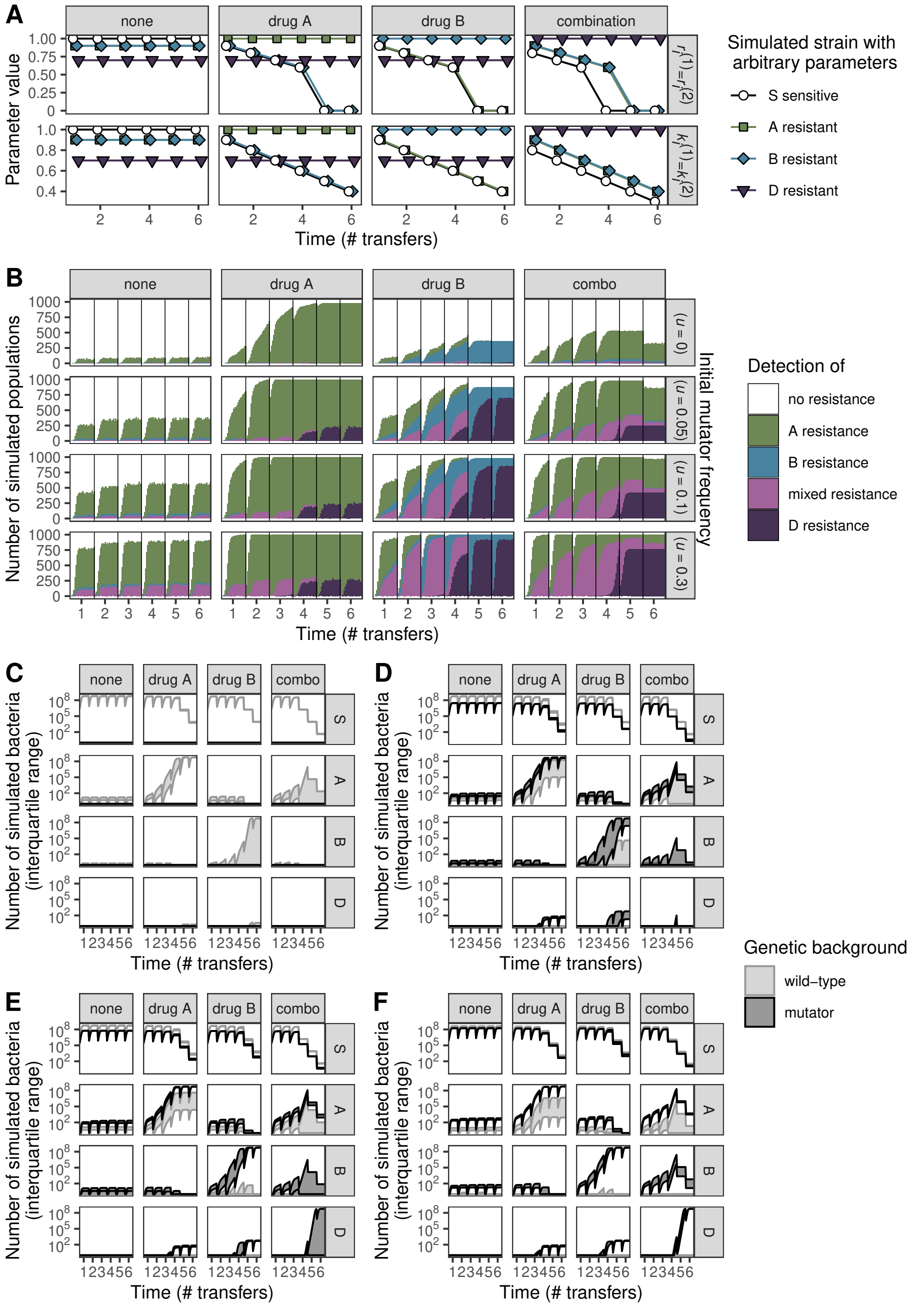


**Supplementary Figure 12:** Simulations with an imposed cost of resistance that also follow a logistic growth curve in pure culture (i.e. Eq. 14, with $r_i^{(1)} = r_i^{(2)}$ and $k_i^{(1)} = k_i^{(2)}$) for $n = 1000$ replicate simulations. A) Parameter values chosen so that simulated strains always exhibit a cost of resistance (points displaced on the horizontal axis for clarity). B) The proportion of populations with each strain type. C–F) Population dynamics of simulated resistance evolution for mutator frequencies ‘none’ (C, $u = 0$), ‘low’ (D, $u = 0.05$), ‘intermediate’ (E, $u = 0.1$), ‘high’ (F, $u = 0.3$).

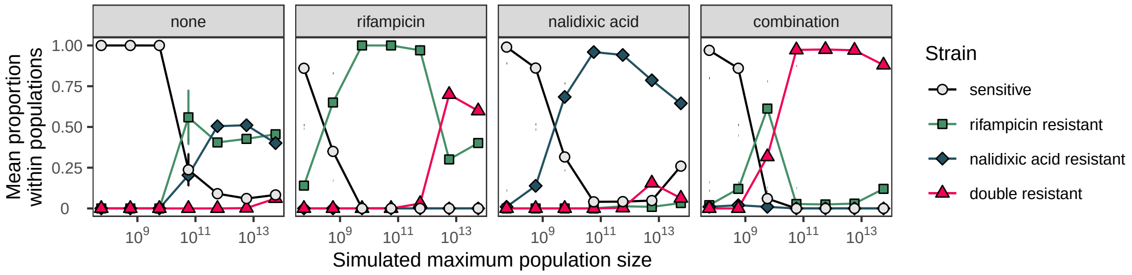


**Supplementary Figure 13:** Simulations of multi-resistance in the absence of mutators indicates multi-resistance can also evolve in very large populations. Proportion of each strain type (±SD, $n = 1000$ replicate simulations) found within simulated populations.

**Supplementary Table 10:** Effects of initial mutator frequency and antibiotic treatment on resistance state observed at the end of the simulations. Coefficients come from fitting a Bayesian categorical regression model to the simulations. Treatment contrasts on the logit scale are shown. (* denotes 95% credible intervals excluding zero).

| Resistance state | Coefficient | Estimate | 95% credible interval | |
|---|---|---|---|---|
| rifampicin resistance | intercept | -1.08 | (-1.58, -0.60) | * |
| | low | 1.41 | (0.87, 1.96) | * |
| | intermediate | 2.09 | (1.50, 2.70) | * |
| | high | 3.66 | (2.57, 4.97) | * |
| | rifampicin | 2.03 | (1.34, 2.71) | * |
| | nalidixic acid | -0.97 | (-1.56, -0.40) | * |
| | combination | -1.34 | (-1.99, -0.71) | * |
| nalidixic acid resistance | intercept | -2.89 | (-3.80, -2.06) | * |
| | low | 0.81 | (-0.04, 1.62) | |
| | intermediate | 0.90 | (-0.10, 1.85) | |
| | high | 2.45 | (0.84, 4.00) | * |
| | rifampicin | -2.94 | (-8.42, 0.38) | |
| | nalidixic acid | 1.29 | (0.45, 2.19) | * |
| | combination | -0.15 | (-1.31, 0.91) | |
| mixed resistance | intercept | -3.37 | (-4.89, -2.17) | * |
| | low | 1.56 | (-0.02, 3.30) | |
| | intermediate | 3.30 | (1.95, 4.89) | * |
| | high | 6.14 | (4.55, 8.07) | * |
| | rifampicin | -3.26 | (-8.59, -0.12) | * |
| | nalidixic acid | -2.58 | (-4.21, -1.21) | * |
| | combination | -2.37 | (-4.01, -1.04) | * |
| double resistance | intercept | -11.34 | (-14.33, -9.15) | * |
| | low | 6.43 | (4.81, 8.75) | * |
| | intermediate | 7.38 | (5.75, 9.72) | * |
| | high | 10.44 | (8.50, 13.01) | * |
| | rifampicin | 8.36 | (6.66, 10.50) | * |
| | nalidixic acid | 5.31 | (3.72, 7.36) | * |
| | combination | 5.72 | (4.12, 7.81) | * |

### Supplementary Information Part 4

### Additional Supplementary Figures

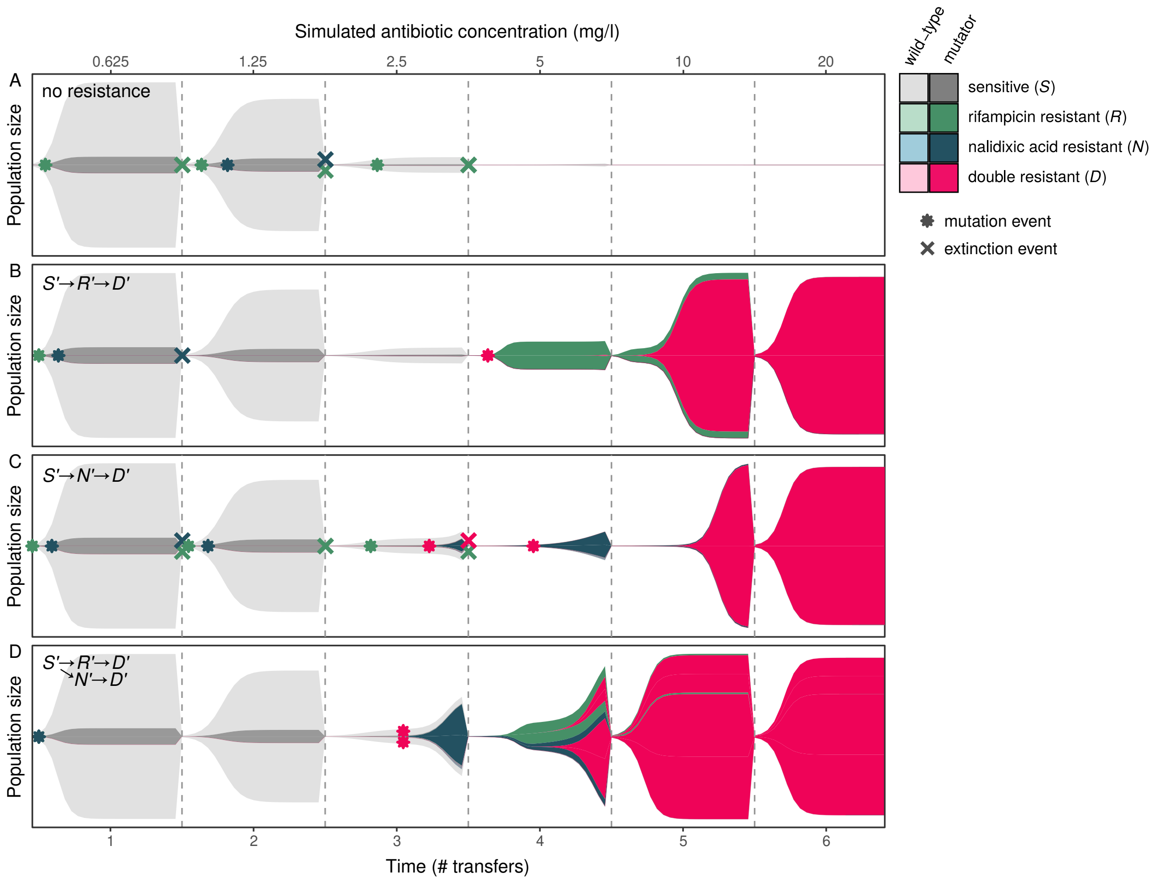


**Supplementary Figure 14: Examples of the four most common types of within-population progression from sensitive to multi-resistant in the combination treatment, demonstrating the emergence of resistance via sequential acquisition of single-drug resistance mutations.** Dynamics are described as follows: A. Single resistance emerged, but failed to establish and ultimately no double resistance is observed. B. Rifampicin resistance establishes first, followed by double resistance (equivalent to main text Figure 4A). C. Nalidixic acid resistance establishes first, followed by double resistance. D. Rifampicin resistance and nalidixic acid resistance both established, followed by double resistance arising in both genetic backgrounds. Areas correspond to the population size of each type. Examples shown are individual replicates from the 'intermediate' initial mutator frequency ($u = 0.1$) treatment.

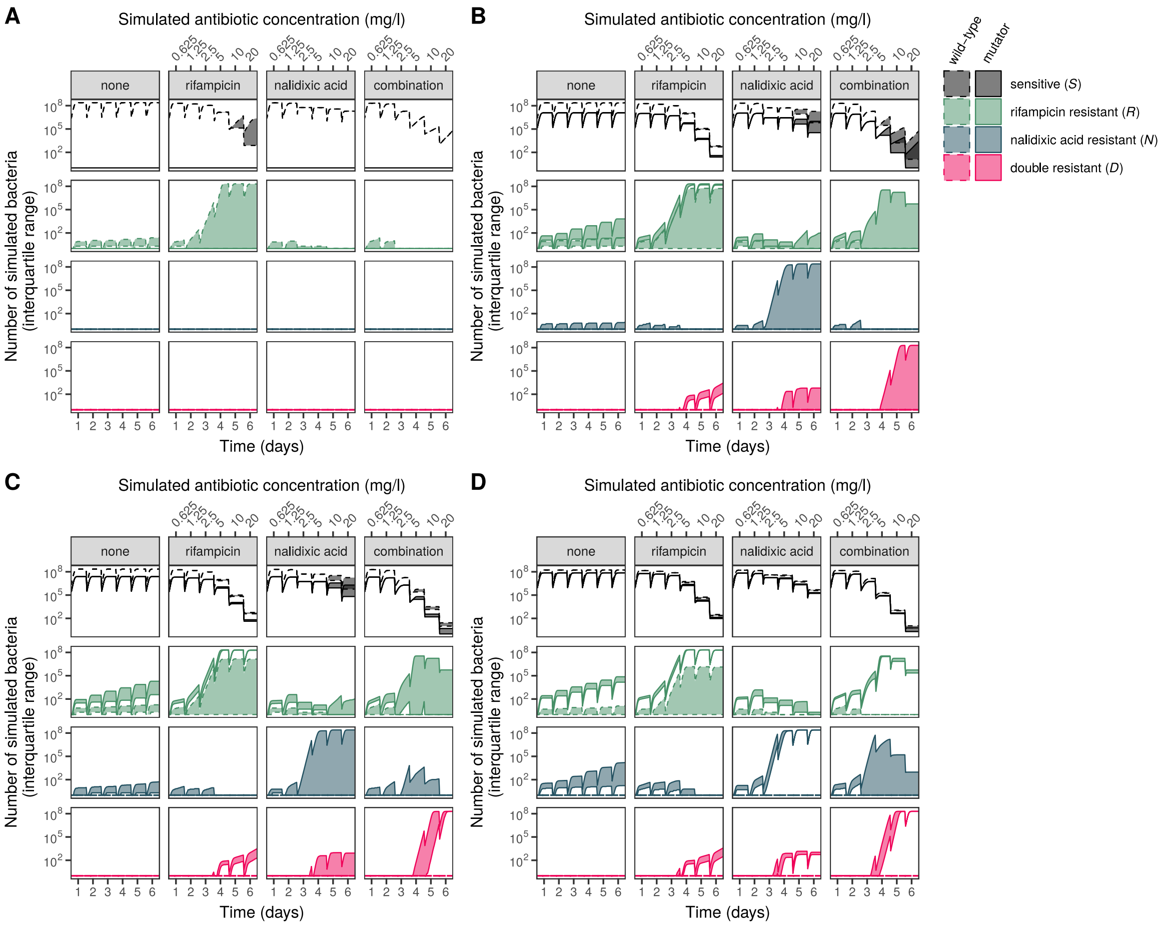


**Supplementary Figure 15: Population dynamics of simulated resistance evolution, demonstrating that multi-resistance swept toward fixation only in the simulated combination treatment.** Interquartile range (IQR) of the number of bacteria of each resistance type over time for the four simulated treatments for $n = 1000$ replicate simulations. Areas indicate the interquartile range (25% and 75% quantiles) of the numbers of bacteria of each resistance type from $n = 1000$ replicate stochastic simulations. Panels A–D show different initial mutator frequencies: 'none' ($u = 0$), 'low' ($u = 0.05$), 'intermediate' ($u = 0.1$, as shown in main text Figure 4), 'high' ($u = 0.3$).
